# Distinct Roles of Central and Peripheral Vision in Rapid Scene Understanding

**DOI:** 10.1101/2025.07.21.665750

**Authors:** Byron A. Johnson, Ansh K. Soni, Shravan Murlidaran, Michael Beyeler, Miguel P. Eckstein

## Abstract

Central and peripheral vision loss, caused by conditions such as age-related macular degeneration and retinitis pigmentosa, disrupt visual processing in distinct ways, yet their impact on natural scene perception remains poorly understood. Here, we used a real-time, gaze-contingent simulation to examine how central vision loss and peripheral vision loss alter eye movements and scene understanding. Sighted participants (*n* = 32, 5 males) viewed 120 natural scenes under one- or three-saccade constraints and described each scene; description quality was quantified via semantic similarity to ground-truth responses. Peripheral vision loss observers produced significantly less informative descriptions than both central vision loss and control participants, particularly for social interaction scenes, suggesting that peripheral vision is critical for rapid extraction of scene semantics. In contrast, central vision loss primarily disrupted oculomotor behavior, including increased saccade amplitudes, delayed saccade initiation, and reduced inter-subject fixation consistency. Description quality was not predicted by fixation similarity to controls, but by fixations to annotated humans and critical objects, underscoring the role of semantically informative sampling. These results reveal a dissociation between perceptual and oculomotor consequences of vision loss and highlight the neural importance of peripheral input for natural scene understanding.

**Significance statement:** Understanding how vision loss affects real-world perception requires disentangling the distinct contributions of central and peripheral vision. Using gaze-contingent simulations of biologically plausible scotomas, we show that peripheral vision loss degrades scene understanding more severely than central loss, particularly for socially meaningful content. In contrast, central vision loss led to more pronounced changes in eye movement planning, including larger and delayed saccades. These findings reveal a dissociation between perceptual and oculomotor consequences of visual field loss, suggesting that peripheral input plays a critical role in rapid semantic processing of natural scenes. Our results underscore the importance of evaluating functional vision using ecologically valid tasks and could inform future strategies in low vision rehabilitation.

## 1 Introduction

Chronic vision loss alters both perceptual processing and eye movement behavior, yet its impact on real-world scene understanding remains poorly understood. Two leading causes of progressive blindness (age-related macular degeneration and retinitis pigmentosa) produce bilateral scotomas that impair vision either centrally (central vision loss (CVL)) or peripherally (peripheral vision loss (PVL)) (Verghese et al., 2021; Shintani et al., 2009). These conditions disrupt the typical mapping between visual input and retinal location, often leading to altered gaze strategies (Legge and Chung, 2016; Guadron et al., 2023; Janssen and Verghese, 2016; Vullings et al., 2022; Seiple et al., 2013).

Previous studies have assessed the perceptual consequences of low vision using psychophysical tasks such as reading, face identification, and visual search (McIlreavy et al., 2012; Geringswald and Pollmann, 2015; Vice et al., 2022). Others have examined scene categorization (or “gist”) using real-world images for patients with CVL or PVL (Tran et al., 2010; Thibaut et al., 2014; Peyrin et al., 2017). Yet these tasks capture only coarse semantic judgments and fail to reveal how vision loss affects higher-level understanding of real-world content. Richer scene interpretation, such as describing social interactions, requires extracting object relationships, context, and meaning beyond simple identification (Greene et al., 2015).

Only a handful of studies have approached this question directly. Costela and colleagues found that patients with CVL gave less consistent verbal descriptions of dynamic scenes than sighted controls, suggesting degraded scene understanding (Costela et al., 2019). However, their study lacked eye-tracking, limiting insight into how visual sampling may have contributed. More generally, the link between gaze behavior and semantic scene interpretation in vision loss remains unclear.

Gaze-contingent displays offer a powerful tool to simulate central or peripheral vision loss in a rapid and controlled manner (Kasowski et al., 2023). These paradigms preserve naturalistic viewing while selectively removing information from specific retinal regions. Simulations of CVL have been shown to affect face perception, reading, and oculomotor strategies (Tsank and Eckstein, 2017; Yu and Kwon, 2023; Vice et al., 2022), with consistent findings of increased saccade amplitude and delayed search performance (Nuthmann, 2014; McIlreavy et al., 2012). However, no prior study has applied these methods to open-ended scene understanding tasks or directly contrasted the effects of central and peripheral loss on semantic interpretation.

To fill this gap, we combined biologically plausible gaze-contingent scotoma simulations with a rapid scene description task involving real-world images. Participants viewed scenes for one or three saccades under simulated CVL, PVL, or control conditions, and were asked to describe the scene content. Eye movements were recorded throughout, allowing us to test how visual sampling, fixation patterns, and semantic understanding are affected by different types of visual field loss. Our approach bridges the gap between low-level gaze metrics and high-level scene comprehension, offering new insights into how central and peripheral vision support everyday perception.

## 2 Methods

### 2.1 Participants

We recruited 32 undergraduate students (5 male, 27 female, mean age of 19.88 years old). Participants were recruited from the Psychological and Brain Sciences Department at the University of California, Santa Barbara and completed the study for course credit. All participants provided informed consent. Experimental protocols were approved in accordance with the Institutional Review Board (IRB) of the University of California, Santa Barbara. All participants had normal or corrected-to-normal vision. None of them reported having any known visual impairments. Any potential subject that could not pass the initial calibration test did not advance to the first trial and was excluded from the experiment.

### 2.2 Apparatus

Python was used to program and run the experiment with scripts developed in-house. We specifically used PsychoPy (Peirce et al., 2019) to present visual stimuli and control the eye tracker. The monitor used was an Alienware AW2524H monitor with 1280 × 1024 pixel resolution (31.3 × 24.8^◦^) and a refresh speed of 480 Hz. The experiment was run on a Windows PC with an AMD Ryzen 7 3700X Processor (3.60 GHz). Participants were seated 68.6 cm away from the monitor with a headrest and chair to minimize head movements.

Participants’ eyes were tracked monocularly using a SR Research Eyelink 1000 plus Tower Mount (SR Research ltd., Ontario, Canada) eye tracker. If calibration was unsuccessful with the right eye for whatever reason, we recorded from the left eye for that participant for the entirety of the experiment. A velocity threshold of 30 ^◦^ s^−1^ and acceleration threshold of 9500 ^◦^ s^−2^ was used to detect saccade events. The sampling rate for the eye tracker was 2000 Hz. Participants performed nine point calibration before each block. If two or more broken fixations occurred during the forced fixation portion of each trial, participants were re - calibrated.

### 2.3 Stimuli

Participants completed a rapid scene understanding task. Each image was a real-world scene scaled to 881.2 ± 61.7 × 600 pixels (21.8 × 14.7^◦^). A total of 120 real-world scenes were used:

- “Social interaction” scenes: Half of the scenes were from a novel dataset of photos collected at the University of California, Santa Barbara. In this dataset, a premise for each scene was generated based on one or more actors interacting with one or more objects or people. Photographs of people acting as characters for these roles were taken to generate a dataset of social interaction photos. These scenes could also contain actors in the background. Example “social interaction” scenes are shown in the top row of Figure 1.
- “Neutral” scenes: The other half of the scenes were pseudo-randomly selected from the publicly available Microsoft Common Objects in Context (MSCOCO) image data set (Lin et al., 2014) used for machine learning and object recognition tasks. While half of the MSCOCO images contained at least one person (*n* = 30), others only contained empty rooms or animals. Importantly, the MSCOCO dataset was specifically designed for benchmarking computer vision model performance for object recognition. The bottom row of Figure 1 shows example “neutral” scenes.

**Figure 1.**
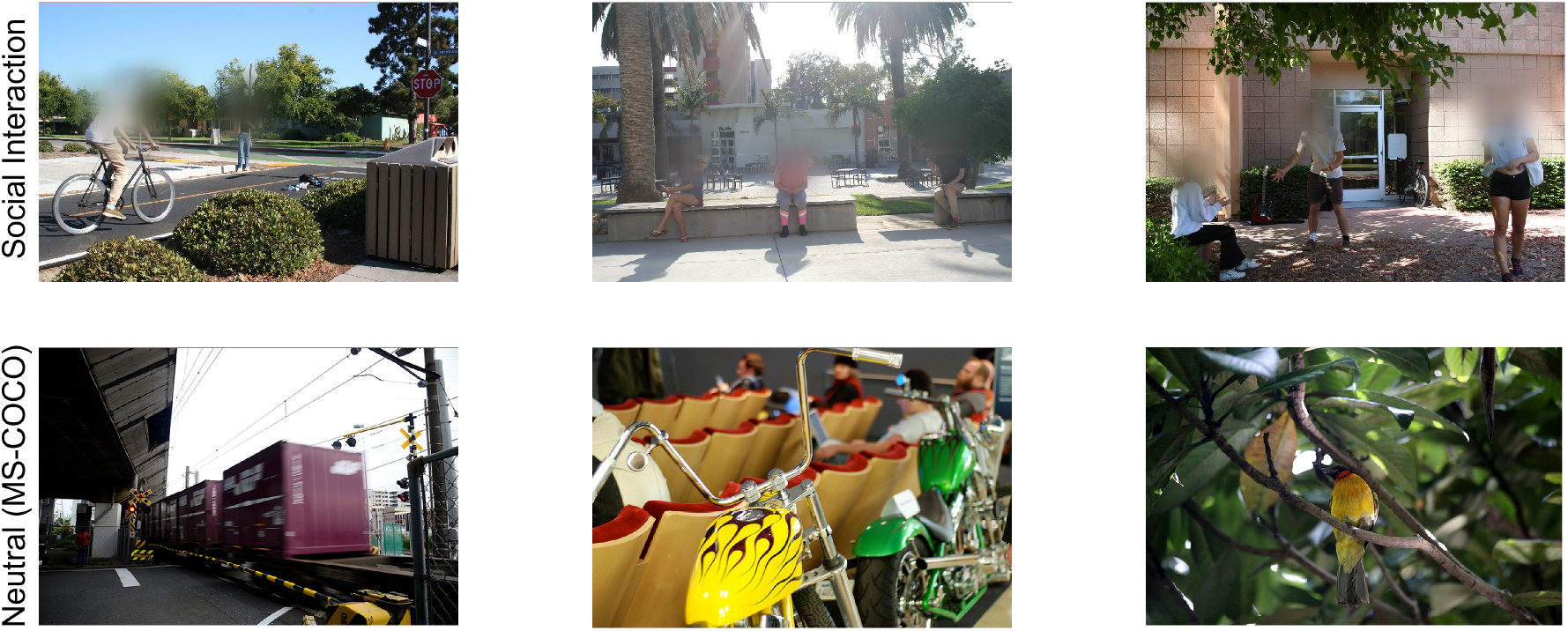
Representative images from the two scene categories used in the study. The top row shows social interaction scenes characterized by human presence and observable social behavior. The bottom row includes neutral scenes from the MS-COCO dataset, containing inanimate objects or animals without overt human interaction. Faces in example social interaction scenes are blurred for sharing of this manuscript.

To explore the effects of low vision viewing conditions on scene understanding, participants were assigned to one of three conditions: none (control, *n* = 10), PVL (*n* = 11), and CVL (*n* = 11). One participant from the control group was unable to finish the study so they were excluded from the dataset. Participants viewed every scene in their assigned condition (e.g. between - subjects design). Participants were not aware of the experimental hypotheses and were not explicitly told the details of their viewing condition.

To simulate PVL, concentric blur was applied to the image leaving a clear window with a diameter of 10^◦^. While previous simulations have used a gray mask (Yu and Kwon, 2023; Vice et al., 2022), we used Gaussian blur from OpenCV (Bradski, 2000) for the simulated scotomas. This is because previous work with patients has shown that they are not always aware of the presence or location of their scotoma (Hartong et al., 2006; Fletcher et al., 2012; Peli et al., 2023). A solid filled mask has sharper edges that are more salient against a blank background, while Gaussian blur can make it more difficult for participants to detect the edges of the scotoma layered on a real-world scene. For CVL, an inner circular patch was blurred with the same Gaussian blur used for PVL. The size of the central scotoma matched the size of the clear window for the PVL condition (10^◦^). This size was based on previous work simulating CVL for a visual search task (Kwon et al., 2013). Figure 2 shows an example scene of all three viewing conditions.

**Figure 2.**
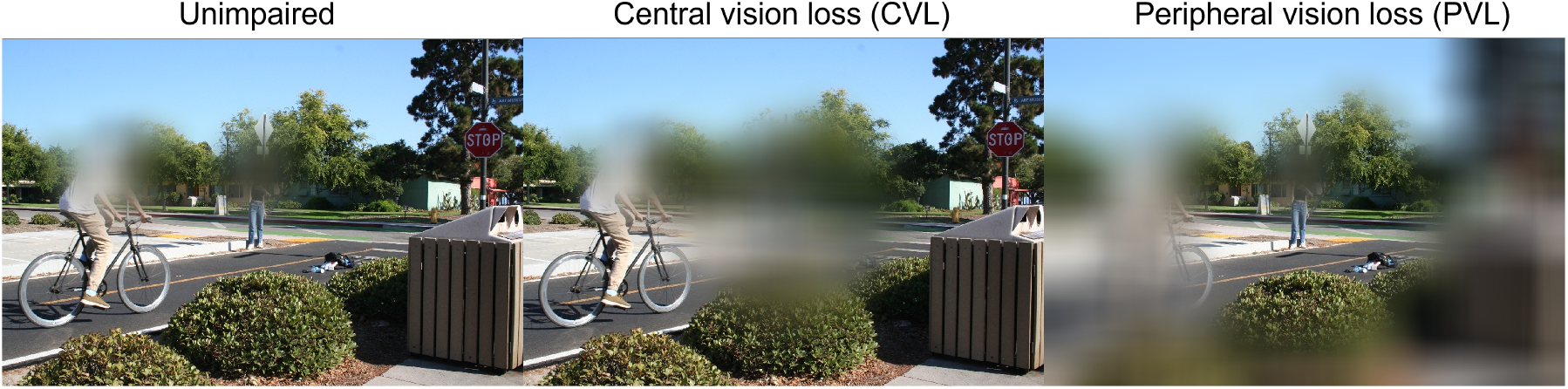
Simulated viewing conditions for a representative social interaction scene. The left panel shows the original image as seen under unimpaired vision. The center panel simulates central vision loss (CVL), with a central scotoma spanning 10^◦^ of visual angle, obscuring the person and backpack on the right. The right panel simulates peripheral vision loss (PVL), with a 10^◦^ clear central window, obscuring the approaching biker on the left. The full scene subtends 21.8 × 14.7^◦^ of visual angle. Faces in example social interaction scenes are blurred for sharing of this manuscript.

The display updated based on real-time recordings from the eye tracker to make the PVL and CVL simulations gaze-contingent. Before each trial, the stimulus was processed with the scotoma filtered at different locations of the image by subsampling every 40 pixels (1^◦^) of the scene in order to present the simulation with minimal lag. Once the scene appeared, the center of the clear window or central scotoma appeared on the screen at the participants current fixation location and moved as participants made eye movements. The average delay between the eye tracker and the experiment monitor was 6.89 ± 1.21 ms (see Figure S1.)

### 2.4 Procedure

#### 2.4.1 Scene Presentation

An overview of the experiment can be seen in Figure 3. The experiment was broken up into 12 blocks with 10 trials each, resulting in 120 trials. Before the experiment, scenes were randomly assigned to each block. While every participant had the same block sequence, the order of scenes within the block was randomized for each participant. The experiment was programmed to close at the end of each block. Participants were allowed to take as many breaks as they requested throughout the experiment. Experimental sessions were one to two hours in length and took place over two or more days.

**Figure 3.**
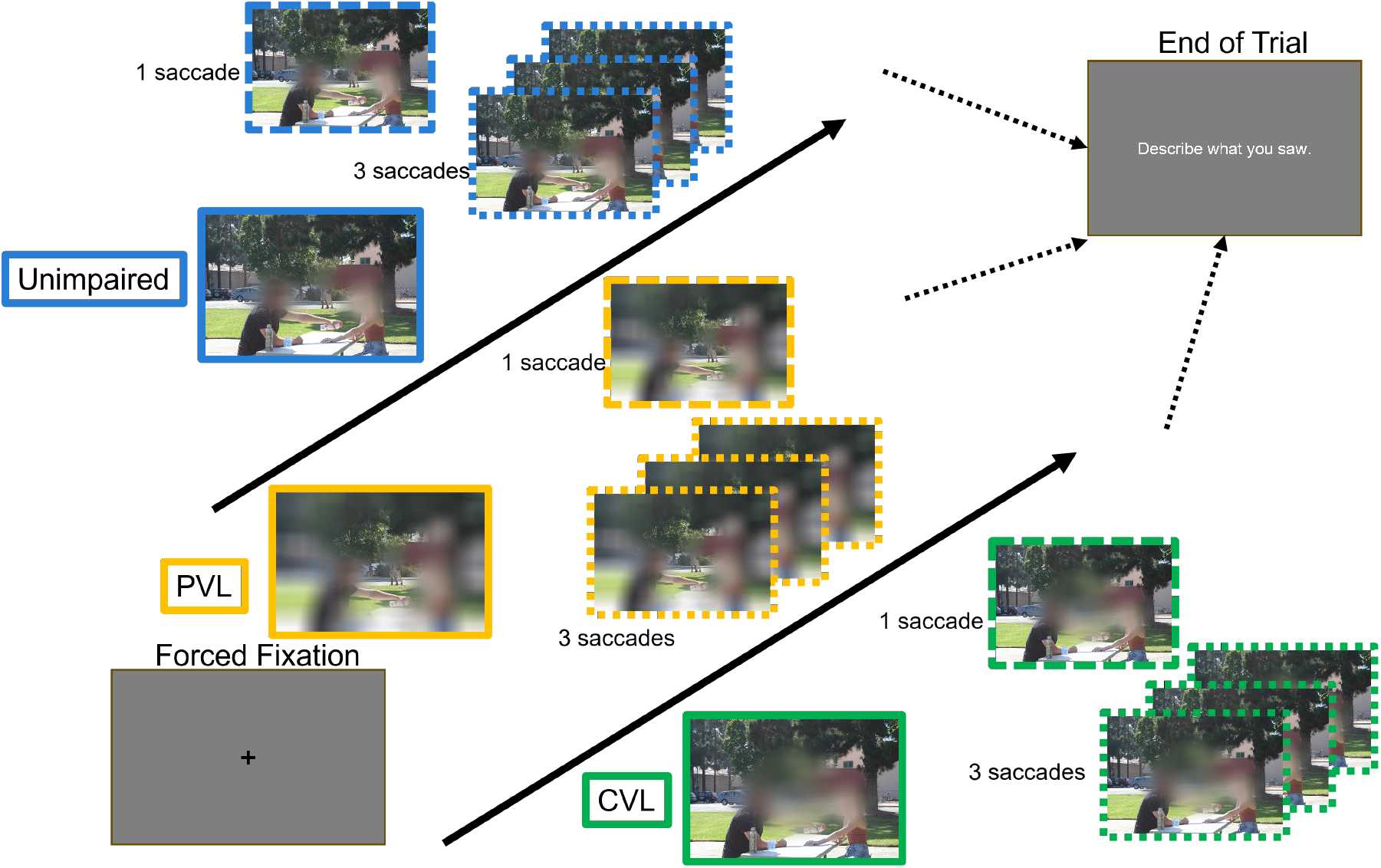
Overview of the experimental procedure. Each trial began with a central fixation cross. Participants (n = 32) viewed a natural scene under one of three simulated viewing conditions: unimpaired, peripheral vision loss (PVL), or central vision loss (CVL). Depending on the trial, participants were allowed either one or three saccades before the scene disappeared (randomized). They were then prompted to describe what they saw. No feedback was given. The same set of scenes was used across conditions and saccade limits, with presentation order counterbalanced across participants. Faces in example social interaction scenes are blurred for sharing of this manuscript.

Participants completed a rapid scene understanding task viewing all stimuli in their assigned simulation condition. After successful calibration, participants were shown a fixation cross in the middle of the screen. To start the trial, participants had to press the ‘space’ key while looking at the fixation cross for a random amount of time selected from a uniform distribution between 250 and 750 ms. If the participant moved their eyes away from the fixation cross for any reason, a “broken fixation” message would appear. Participants were re-calibrated after frequent broken fixations. Participant data was automatically saved after each trial. If the participant had to take a break or re-calibrate between trials, they would exit the program. Upon restarting, they would re-calibrate and continue with the next trial in sequence.

If the participant fixated the central cross successfully (250 to 750 ms), the scene would appear. The amount of time for the stimulus presentation was dependent on the number of saccades made by the participant while viewing the scene. Before the experiment, half of the scenes were randomly assigned to allow for one saccade or three saccades. If a scene was assigned to allow for one saccade, upon stimulus onset, participants could make one saccade starting from the initial central fixation upon scene onset to their second fixation. If a scene was assigned to allow for three saccades, participants could make four fixations (initial central fixation at scene onset until completion of the fourth fixation). The simulated scotoma would update based on the participant’s current fixation location. Once the saccade limit was reached, upon the start of the next saccade, the scene disappeared and a gray screen was shown. If the saccade limit was not reached, the total maximum scene presentation time was 10 s. This fast presentation for scene perception is referred to as “rapid scene understanding”.

#### 2.4.2 Response

After the scene disappeared, participants were instructed to type a description of the scene in English. There was no strict length limit to their response but participants were encouraged to keep their descriptions shorter than four sentences. Text would appear on the screen as participants typed. No other feedback was provided about the quality of their descriptions.

Verbal instructions for the scene description were given at the beginning of the experiment.

The following instructions were presented on screen at the beginning of the experiment and every block:

1. Double check descriptions for errors before submitting.
2. Use complete sentences only. No one word sentences.
3. Do not use proper nouns or names.
4. Gender and pronouns can be used.
5. Start your description with “There was a…”
6. Only describe what you saw. Do not put “unclear” or “could not see it”.

### 2.5 Metrics

#### 2.5.1 Scene Descriptions

The semantic similarity between a ground-truth description of the scene and each participant response was computed as follows:

- **Generating ground truth descriptions of each scene:** To generate ground truth descriptions of each scene, a different group of five participants (5 female, mean age of 20.2 years old) were given the same scene task with unlimited viewing time and no simulated impairment. This provided five sets of ground truth descriptions to be used to evaluate the semantic similarity of responses from the rapid scene understanding task. To evaluate the upper limits of the ground truth descriptions, a separate group of five participants (5 female, mean age of 19.8 years old) compared four sets of ground truth descriptions for all scenes to one set of ground truth descriptions. One ground truth description set was duplicated, randomly assigned to scenes, and also included to test the lower limit of the ground truth descriptions.
- **Rating participant descriptions:** Two measures were used to evaluate the quality of participant descriptions.

– The first measure is based on **human ratings** of descriptions and was inspired by work that examined how humans judge the similarity of pairs of words and text (Wang et al., 2023; Nguyen et al., 2014). A third group of four participants (1 male, mean age of 20 years old) compared every response to the ground truth description for each scene. Every response was rated on a scale of 1 to 10 where 1 meant that the response did not semantically match the ground truth description and 10 meant that the response semantically matched the ground truth description. Each rater (*n* = 4) provided a score for every participant response set (*n* = 32) for all scenes (*n* = 120), totaling 15,360 ratings. The average rating between the four raters provided 3,840 measures of semantic similarity to the ground truth descriptions. Human ratings were generated from the descriptions only (the scene image was not provided to the raters). Importantly, only one ground truth description set was used for the human rating analyses. To determine variability in ratings, inter-rater reliability and Pearson’s correlations were calculated between raters. Inter-rater reliability calculates Cohen’s Kappa between each rater to determine the level of agreement on categorical data (scores 1 - 10) as:

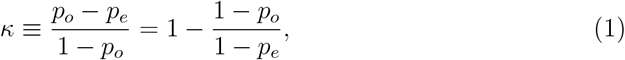

where *p*_*o*_ is the agreement between two raters, *p*_*e*_ is hypothetical chance agreement between the 10 scores, resulting in *κ* ∈ [0, 1] with 0 meaning no agreement as predicted by chance and 1 meaning complete agreement.
– The second measure for analyzing participant responses generates embeddings based on **Chat Generative Pre-Trained Transformer (GPT)-4** (Open AI, Inc., California, United States). GPT-4 can be used to compare text in an embedding space. These embeddings can be used to compute **cosine similarities** between pairs of ground truth descriptions and participant responses for one scene. GPT-4 is popular for its impressive language inference abilities based on Large Language Models (LLMs) and can be used to quantify potential differences in participant responses. We used GPT-4 to generate scores of semantic similarity between ground truth descriptions as candidate text and participant responses as reference sentences:

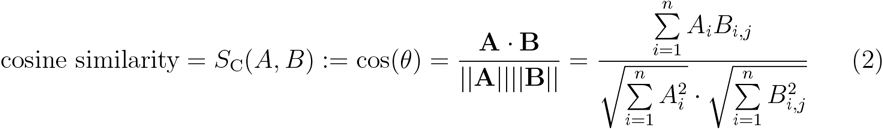

where for each scene *i* and each participant response *j*, the candidate ground truth description *A*_*i*_ is paired with the participant response for that scene *B*_*i,j*_. The dot product of the components of each sentence are normalized by the magnitude of both descriptions combined in Euclidean space, providing the cosine similarity score. This value can range from -1 to 1 with 0 meaning weakly related and 1 meaning exactly the same.

#### 2.5.2 Eye Movement Metrics

Saccade amplitude is the third measure and was calculated as the distance between the start of a saccade (end of a fixation) and the end of a saccade (start of a fixation). The fourth measure is saccade latency and was defined as the total amount of time between the scene appearing and the participant making a saccade. This was calculated for every trial and every recorded eye movement. If more than one saccade was allowed to view the scene, the latency was measured as the time between the end of the previous saccade and the beginning of the next saccade (i.e. fixation dwell time).

#### 2.5.3 Fixation Heat Maps

Analysis of participant eye movement behavior was based on five metrics to capture the spatial and temporal features of saccades. Distributions of participant fixation locations for a scene were measured by transforming fixation locations for every scene into separate heat maps for each viewing condition. To assess how fixations varied across viewing conditions (controls, PVL, and CVL) we compared fixation heat map correlations across viewing conditions to groups of different subjects in the control condition. Each heat map was generated by randomly sampling subjects (*n* = 4) into four groups for testing: one group of controls (“control comparability group”), a different group of controls to compare to the control comparability group, a group of PVL, and a group of CVL. All fixations recorded during the stimulus presentation were convolved with a Gaussian filter (39 × 39 pixels, or approximately 1 × 1^◦^), providing a spatial representation of where participants fixated during each trial. Each heat map has a color map representing fixation density, where red indicates a high number of fixations relative to the rest of the map. Correlations between each control comparability group heat map and each viewing condition fixation heat map were computed for every scene.

#### 2.5.4 Scene Segmentation and Frequency of Fixations to Objects

Since we used real-world scenes containing people and objects, we wanted to understand if fixations to specific objects were affected by the viewing condition. The second measure based on eye movement behavior used annotations to determine how often participants fixated objects. One annotator (1 male, 19 years old) had unlimited time to outline faces and bodies of humans and animals as well as critical objects for all scenes. The annotations were created using Make Sense (Skalski, 2019) and saved into a .JSON file. Fixation locations separated by viewing condition and annotations were layered onto each scene, then the number of fixations that were located within an annotation were counted. Examples of annotated social interaction and neutral scenes are in Figure S7.

Given that 61 of the 120 scenes presented (45 social interaction and 16 neutral) contained people located in the periphery (greater than 5 degrees visual angle) with respect to the fovea at the start of the trial) we classified each scene (ad hoc) with a binary label to determine if the effects of viewing condition on descriptions were affected by peripheral arrangements of people. Figure S4 shows an example scene classified as having a person located in the periphery upon stimulus onset. We repeated this labeling process for critical objects in the scenes. The critical object was defined as the most important object (or group of objects) needed to accurately describe the scene. Thirty-three of 120 scenes (21 social interaction, 12 neutral) were labeled as either including a critical object that was located in the periphery or a critical object that was located in the center of the scene.

### 2.6 Statistical Analyses

#### 2.6.1 Description Ratings

A linear mixed model lme4 (Bates et al., 2015) was used to assess the variance in human description ratings, including potential interactions between viewing condition (none, PVL, CVL), scene type (social interaction or neutral), and number of saccades allowed (one or three). Participants’ unique identifiers were included as a random factor to account for variability in semantic similarity across subjects. Post hoc comparisons were performed using Šidák correction to test for pairwise differences between conditions.

#### 2.6.2 Analysis of Eye Movement Metrics

For each trial, participants’ first saccade amplitude and first saccade latency were entered into an *F* -test to assess whether these basic eye movement metrics varied across viewing conditions.

#### 2.6.3 Fixation Heat Map Similarity

To assess how fixations varied across viewing conditions, fixation heat maps were generated for each group of subjects. Participants were randomly assigned to groups of four from each viewing condition: control, PVL, CVL, and a separate “control comparability group” composed of different control participants. Fixation locations were convolved with a Gaussian filter to produce a heat map for each scene and group. For each iteration, we computed the correlation between the heat map of an experimental group and the control comparability group. This procedure was repeated 100 times per scene, and average correlation scores were used in a linear mixed model to test whether heat map similarity was influenced by viewing condition, scene type, or number of saccades allowed. Scene ID was included as a random factor to account for variability due to specific scene content. Multiple comparisons were corrected using the Šidák method. A secondary analysis tested whether heat map similarity differed based on whether people or critical objects were located in the periphery.

#### 2.6.4 Fixations to Annotated Objects

The number of fixations to annotated humans and critical objects was computed per scene and viewing condition. A linear mixed model was used to test whether the number of object-directed fixations varied as a function of viewing condition, scene type, or number of saccades allowed. Scene ID was included as a random factor. Post-hoc Šidák-corrected comparisons were performed across conditions, and object arrangement (e.g., peripheral layout) was included as an additional binary predictor.

#### 2.6.5 Relating Semantic Similarity to Eye Movements

To examine the relationship between scene understanding and eye movement behavior, we correlated average human ratings with across-condition fixation heat map correlations on a per-scene basis. This allowed us to assess whether higher description ratings were associated with more control-like fixation patterns. Additional correlations were computed between description ratings and other eye movement metrics to determine how viewing condition modulated behavior across modalities.

## 3. Results

### 3.1 Effects of Simulated Vision Loss on Semantic Scene Understanding

#### 3.1.1 Human Ratings of Semantic Similarity

Figure 4A–B shows how participant descriptions varied by viewing condition (control, PVL, CVL), scene type (social or neutral), and number of saccades allowed (one or three). A linear mixed-effects model was used to predict average human similarity ratings aggregated across four raters for each trial (see Supplemental Material for rating distribution and inter-rater reliability characteristics). Fixed effects included viewing condition, scene type, and saccade number, and random intercepts were included for both participant and scene. The model using unique subject and scene identifiers as random intercepts significantly outperformed a subject intercept-only baseline (*χ*^2^(1) = 1465.95, *p* < .001; conditional *R*^2^ = .37, marginal *R*^2^ = .07).

**Figure 4.**
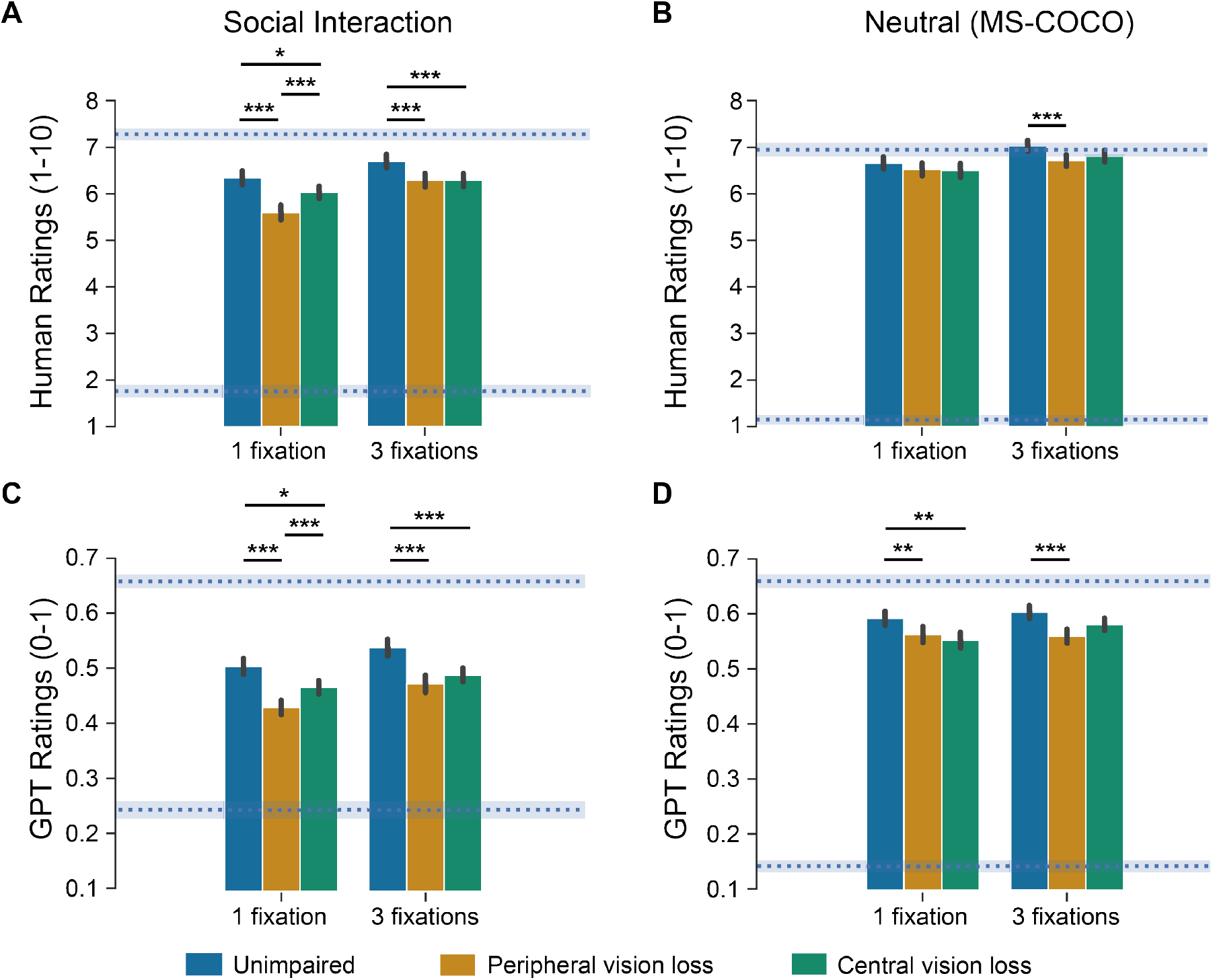
Semantic similarity scores for participant descriptions across viewing conditions, scene types, and number of saccades allowed. (A–B) Mean human ratings of description quality. (C–D) GPT-4 cosine similarity scores comparing participant descriptions to ground truth. For social interaction scenes with one saccade, PVL descriptions were rated significantly lower than both controls and CVL by human raters (A) and received lower GPT-4 scores (C). With three saccades, both PVL and CVL descriptions were rated lower than controls for social scenes. For neutral scenes, PVL descriptions were rated lower than controls by human raters with one saccade (B), and by GPT-4 with either one or three saccades (D). Horizontal dashed blue lines (and shaded areas) denote the range of ground truth scores for each scene type. Human rating ground truths: social = 7.21 ± 0.12 (upper), 1.74 ± 0.12 (lower); neutral = 6.92 ± 0.13 / 1.17 ± 0.09. GPT-4 scores: social = 0.65 ± 0.01 / 0.25 ± 0.01; neutral = 0.65 ± 0.01 / 0.17 ± 0.01.

A three-way ANOVA on the fitted model revealed a significant interaction between viewing condition, scene type, and number of saccades (*F* (2, 3712) = 3.99, *p* = .019), indicating that the quality of scene descriptions depended jointly on visual loss type, content type, and allowed viewing time.

To validate assumptions for the model, we first tested for homogeneity of variance across groups using Levene’s test, which indicated unequal variances (*F* (2, 3837) = 8.83, *p* < .001). A Welch’s ANOVA confirmed a significant difference between at least one pair of group means (*F* (3840) = 35.48, *p* < .001). Inspection of residuals revealed no substantial relationship between fitted values and residual variance, supporting the use of mixed-effects modeling for subsequent analyses.

To interpret the significant three-way interaction, we conducted Šidák-corrected pairwise comparisons. For social scenes with only one saccade allowed, both PVL and CVL groups produced significantly lower similarity ratings than controls (PVL: *M* = −0.52, *p* < .001, *d* = 0.52; CVL: *M* = −0.21, *p* = .02, *d* = 0.22), and PVL ratings were also significantly lower than CVL (*M* = −0.31, *p* < .001, *d* = 0.30). With three saccades, both impaired groups remained significantly lower than controls (PVL: *M* = −0.39, *p* < .001, *d* = 0.39; CVL: *M* = −0.37, *p* < .001, *d* = 0.37), with no difference between PVL and CVL (*M* = 0.03, *p* = 1).

For neutral scenes, the only significant difference emerged when three saccades were allowed: the PVL group rated significantly lower than controls (*M* = −0.28, *p* < .001, *d* = 0.28). No other pairwise differences were significant for neutral scenes (*p* values = .28–1).

#### 3.1.2 LLM-Based Ratings Using GPT-4 Embeddings

To complement the human ratings, we computed semantic similarity scores using GPT-4 sentence embeddings (Figure 4C–D). For each participant description, we computed the cosine similarity to the corresponding ground truth description. These model-based scores were analyzed using the same linear mixed-effects model structure as the human ratings, with fixed effects for viewing condition, scene type, and number of saccades, and random intercepts for participant and scene.

The model revealed a significant three-way interaction between viewing condition, scene type, and number of saccades (*F* (2, 3712) = 4.77, *p* = .009), consistent with the human rating results.

GPT-based similarity scores were significantly correlated with average human ratings across trials (*r* = .73, *p* < .001), indicating convergent validity between the two approaches.

Šidák-corrected pairwise comparisons revealed that, for social scenes with one saccade, both PVL and CVL groups produced significantly lower cosine similarity scores than controls (PVL: *M* = −0.06, *p* < .001, *d* = 0.83; CVL: *M* = −0.02, *p* < .001, *d* = 0.29), with PVL lower than CVL (*M* = −0.04, *p* < .001, *d* = 0.54). When three saccades were allowed, both vision loss groups again differed from controls (PVL: *M* = −0.07, *p* < .001, *d* = 0.92; CVL: *M* = −0.05, *p* < .001, *d* = 0.66), with no difference between PVL and CVL (*M* = 0, *p* = 1).

For neutral scenes with one saccade, both PVL and CVL groups scored lower than controls (PVL: *M* = −0.03, *p* = .001, *d* = 0.38; CVL: *M* = −0.03, *p* = .001, *d* = 0.37), but did not differ from each other (*M* = 0, *p* = 1). With three saccades, only PVL differed significantly from controls (*M* = −0.03, *p* < .001, *d* = 0.39); the difference between PVL and CVL was not significant (*M* = −0.01, *p* = .30).

To test for potential learning effects, we compared human ratings and GPT-based cosine similarities across the first and last experimental blocks (Figure S3). A two-way ANOVA found no effect of block number (*F* (1, 6) = 1.27, *p* = .303), indicating that scene descriptions did not improve over time.

### 3.2 Effects on Eye Movements

Two participants were excluded from the eye movement analyses due to missing data, leaving 30 participants (10 control, 11 PVL, 9 CVL) included in all analyses reported below.

#### 3.2.1 Saccade Amplitude and Latency

To test how simulated vision loss affects the dynamics of early eye movements, we measured the amplitude and latency of the first saccade after scene onset. Prior work suggests that individuals with CVL tend to make varied saccades to compensate for their scotoma (Seiple et al., 2013; Vullings et al., 2022), while those with PVL or CVL may exhibit delayed saccades due to the scotoma (Guadron et al., 2023; Van Der Stigchel et al., 2013).

As shown in Figure 5A, saccade amplitude differed significantly by viewing condition (*F* (2, 3478) = 160.71, *p* < .001). Participants with CVL made larger first saccades than both controls (*M* = −1.06 dva, *p* < .001, *d* = 0.65) and PVL (*M* = −1.20 dva, *p* < .001, *d* = 0.73), with no difference between controls and PVL (*M* = 0.13 dva, *p* = .56).

**Figure 5.**
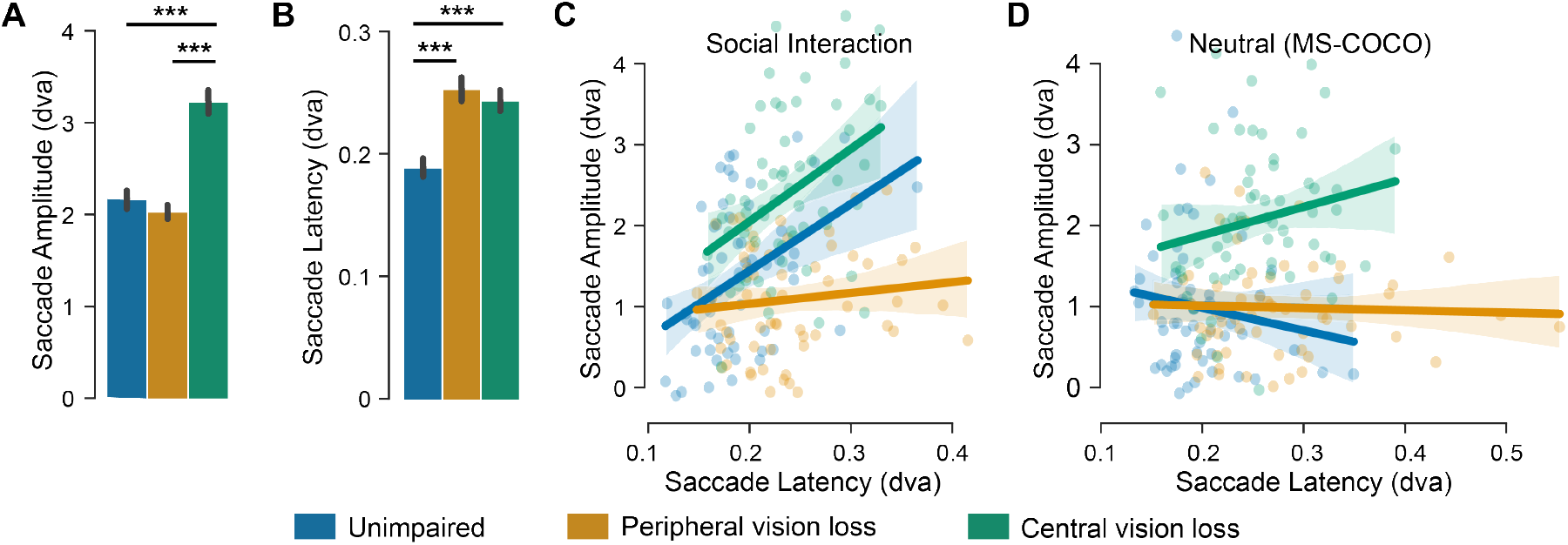
First saccade amplitude and latency. (A) Participants in the CVL condition made significantly larger first saccades than both unimpaired and PVL participants. No significant difference was observed between unimpaired and PVL participants. (B) First saccade latency was significantly longer for both PVL and CVL participants compared to unimpaired controls. There was no significant difference in latency between PVL and CVL conditions. (C–D) Relationship between first saccade amplitude and latency, separated by scene type. A significant correlation was observed for social interaction scenes (C), but not for neutral scenes (D).

Figure 5B shows that saccade latency also varied by condition (*F* (2, 3478) = 125.28, *p* < .001): both PVL and CVL participants were slower than controls (PVL: *M* = +0.06 s, *p* < .001, *d* = 0.71; CVL: *M* = +0.05 s, *p* < .001, *d* = 0.63), with no difference between the impaired groups (*M* = 0.01 s, *p* = .34).

Finally, we assessed whether amplitude and latency were correlated (Figure 5C–D). Across all scenes and participants, these measures were modestly correlated (*r* = .17, *p* = .001), driven primarily by the CVL group (*r* = .26, *p* = .003). The relationship held for social scenes (*r* = .30, *p* < .001) but not neutral scenes (*r* = .09, *p* = .23), and was strongest for controls (*r* = .41, *p* = .001) and CVL participants (*r* = .39, *p* = .001) viewing social scenes.

#### 3.2.2 Fixation Heat Maps

To assess how simulated vision loss alters the spatial distribution of gaze, we compared fixation heat maps between each experimental group and a matched control group. As described in the Methods, heat maps were generated from subsampled participant fixations and correlated with control maps using Pearson’s *r* to quantify spatial similarity (Figure 6A).

**Figure 6.**
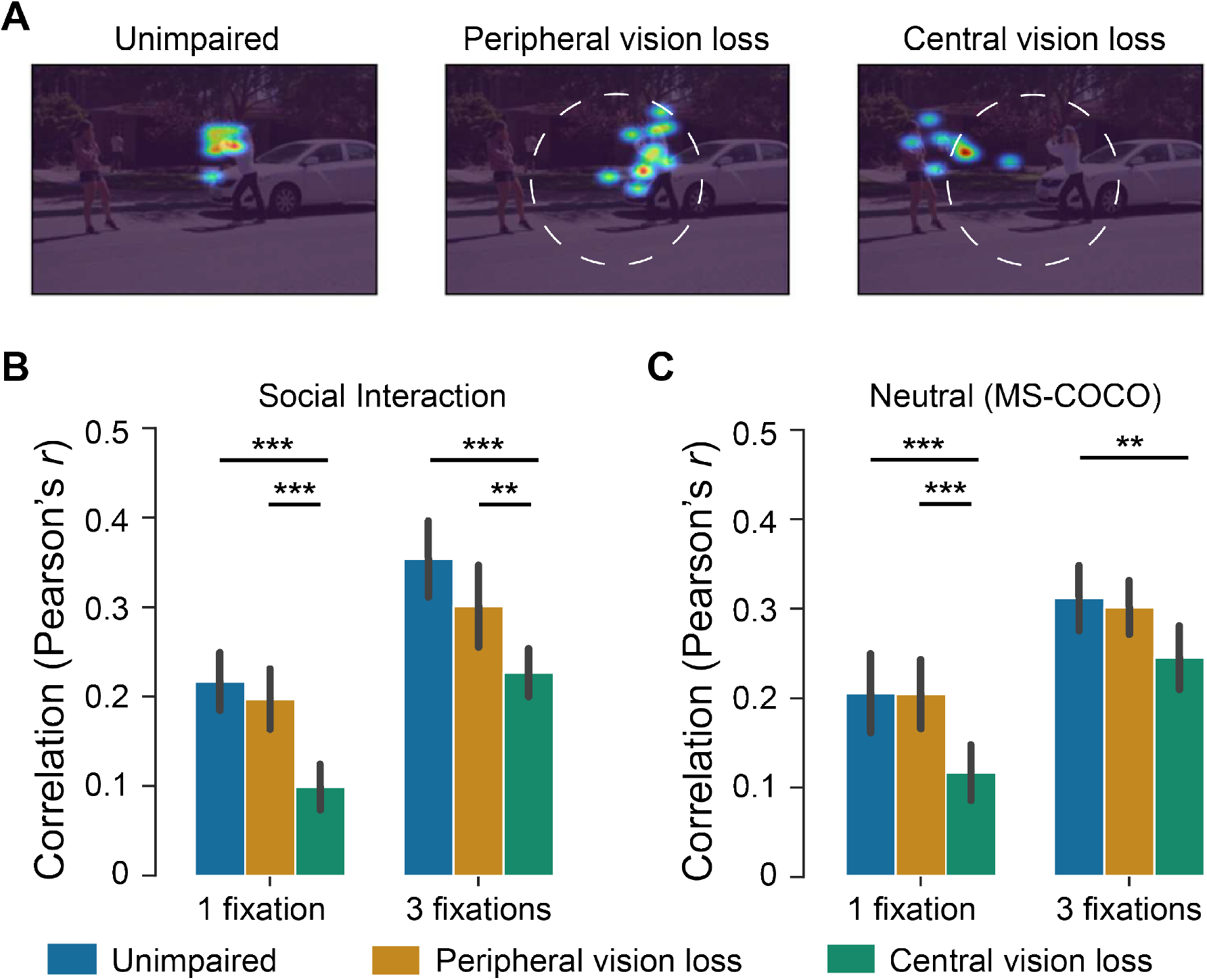
Fixation heat map similarity across viewing conditions. (A) Example fixation heat maps for a social interaction scene, shown for unimpaired (left), PVL (middle), and CVL (right) participants. Heat maps reflect where participants looked during the scene; dashed circles denote the approximate region obscured (CVL) or visible (PVL). Additional heat map examples are shown in Supplemental Figures S4–S6. (B–C) Average Pearson correlation between individual participant heat maps and a control group reference heat map (subsampled, n = 4 participants per condition, repeated 100 times). For both social interaction scenes (B) and neutral MS-COCO scenes (C), heat map similarity was significantly reduced in the CVL group compared to both controls and PVL, across both fixation limits.

Figure 6B–C shows that CVL participants consistently produced less control-like heat maps than either controls or PVL, across both social (Panel B) and neutral scenes (Panel C).

A linear mixed-effect model was fit to determine if heat map correlations differed based on experimental conditions. Viewing condition, saccades allowed, scene type, and their interactions were used as fixed effects. Unique scene identifiers were included as random effects. Maximum likelihood estimation was used and p values were calculated using Satterthwaite’s method of approximation. Scaled residuals for the model were centered around 0 (minimum = −2.42, 1st quantile = −0.6, median = −0.06, 3rd quantile = 0.54, maximum = 2.92) and conditional *R*^2^ = .58 and marginal *R*^2^ = .38. Results of the linear mixed model show a significant difference in heat map correlation between controls and CVL participants (*β* = −0.12, *t*(240) = −6.17, *SE* = 0.02, *p* < .001). The negative *β* coeffiecient suggests that CVL fixation heat maps were less correlated than controls by -.12 on average. The difference in heat map correlations was not signficant between controls and PVL participants (*β* = −0.02, *t*(240) = −1.03, *SE* = 0.02, *p* = .3)

The linear mixed model can only provide results for simple effects differences between controls and PVL or controls and CVL. A second linear mixed model was done to compare differences between PVL with controls and PVL with CVL. This second model revealed a significant difference in heat map correlation between PVL and CVL participants (*β* = -0.1, *t*(240) = −5.146, *SE* = 0.02, *p* < .001), but not between PVL and controls (*β* = 0.02, *t*(240) = 1.02, *SE* = 0.02, *p* = .31) suggesting that CVL fixation heat maps were less correlated than PVL but PVL correlations were not significantly different from controls.

The difference in heat map correlations between one and three saccades allowed was significant (*β* = 0.14, *t*(296.5) = 5.57, *SE* = 0.02, *p* < .001), suggesting that heat maps were 0.14 more correlated when three saccades were allowed instead of one. There was no significant difference in heat map correlations between social interaction and neutral scenes (*β* = −0.01, *t*(296.5) = −0.54, *SE* = 0.02, *p* = .59). The random intercept for the model was significant (*β* = 0.22, *t*(296.5) = 13.16, *SE* = 0.02, *p* < .001) suggesting that heat maps were correlated when fixed effects are ignored. No interaction in the linear mixed model were found to be significant for fixation heat map correlations.

An analysis of variance (ANOVA) was conducted on the linear mixed model to test all possible comparisons between groups. The ANOVA revealed a significant main effect of viewing condition on heat map correlations (*F* (2,232) = 13.91, p *<*.001). There was also a significant main effect of saccades allowed (*F* (1,120) = 81.85, p *<*.001), but not scene type (*F* (1,120) = 0.07, p = .80). The ANOVA did not reveal any significant interactions for the model.

Post hoc comparisons (Šidák-corrected) revealed that CVL heat maps were significantly less correlated with controls than both control and PVL groups across nearly all conditions. For social scenes, heat map similarity was lower with one saccade (vs. control: *M* = 0.12, *p* < .001, *d* = 1.52; vs. PVL: *M* = 0.10, *p* < .001, *d* = 1.27) and three saccades (vs. control: *M* = 0.13, *p* < .001, *d* = 1.64; vs. PVL: *M* = 0.07, *p* < .001, *d* = 1.42). For neutral scenes, CVL groups also differed from both control and PVL with one saccade (control: *M* = 0.09, *p* < .001, *d* = 1.14; PVL: *M* = 0.09, *p* < .001, *d* = 1.13), and from controls with three saccades (*M* = −0.07, *p* = .009, *d* = 0.85), but not significantly from PVL (*M* = 0.06, *p* = .05). No differences were found between control and PVL participants (*p* = .16–1).

ANOVA results for the fixation heat map analysis should be interpreted with caution due to the chance of increased false positives (Type 1 Error) that can occur from the subsampling process. The linear mixed models and ANOVA post hoc results show that CVL participant heat maps were significantly less correlated than controls and PVL participants.

More example heat maps can be found in Appendix Figures S4–S6.

#### 3.2.3 Fixations to Annotated Objects

To evaluate visual access to socially or semantically relevant content, we quantified the number of fixations to annotated humans and critical objects in each scene (see Methods; Figure 7A). As scene type did not significantly interact with any other factors in our model, results are collapsed across social and neutral scenes for clarity (Panels B–C).

**Figure 7.**
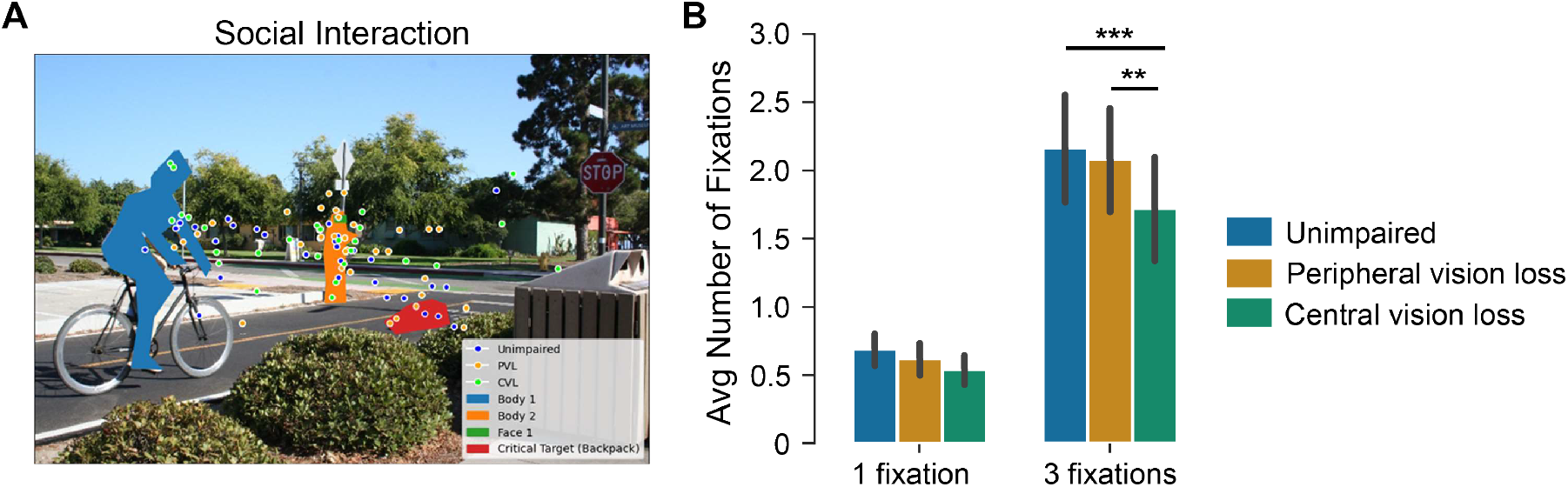
(A) Example scene with annotated humans (orange) and critical task-relevant objects (blue), used to quantify visual access to meaningful scene content (see Methods). (B–C) Average number of fixations to annotated regions by viewing condition and number of saccades allowed. With three saccades, participants in the CVL condition fixated annotated content significantly less often than both controls and PVL participants, regardless of scene type. No significant differences were observed when only one saccade was allowed.

A linear mixed-effects model revealed a significant interaction between viewing condition and number of saccades allowed (*F* (2, 236) = 9.14, *p* < .001). When participants were allowed three saccades (Panel C), those with CVL made significantly fewer fixations to annotated content than both controls (*M* = 0.44, *p* < .001, *d* = 1.31) and PVL participants (*M* = 0.35, *p* < .001, *d* = 1.06). There was no difference between control and PVL groups (*M* = 0.08, *p* = .75), and no significant group differences when only one saccade was allowed (Panel B; *p* values = .17–.88).

### 3.3 Linking Gaze Behavior to Scene Understanding

To assess whether scene understanding was linked to eye movement behavior, we examined correlations between average human description ratings and various eye-tracking metrics across the 360 scene-level data points (i.e., averaged across 32 participants, grouped by scene and viewing condition).

As shown in Figure 8A, there was no significant correlation between average human ratings and global fixation heat map similarity (*r* = .09, *p* = .06), suggesting that the overall spatial distribution of fixations was not predictive of scene comprehension. In contrast, higher description ratings were significantly associated with increased fixations to annotated humans and objects (*r* = .21, *p* < .001; Figure 8B). This relationship held across all viewing conditions (controls: *r* = .26, *p* = .004; PVL: *r* = .20, *p* = .03; CVL: *r* = .18, *p* = .04), suggesting that the informativeness of participant descriptions was better predicted by socially or semantically relevant content than by the overall shape of their gaze distribution.

**Figure 8.**
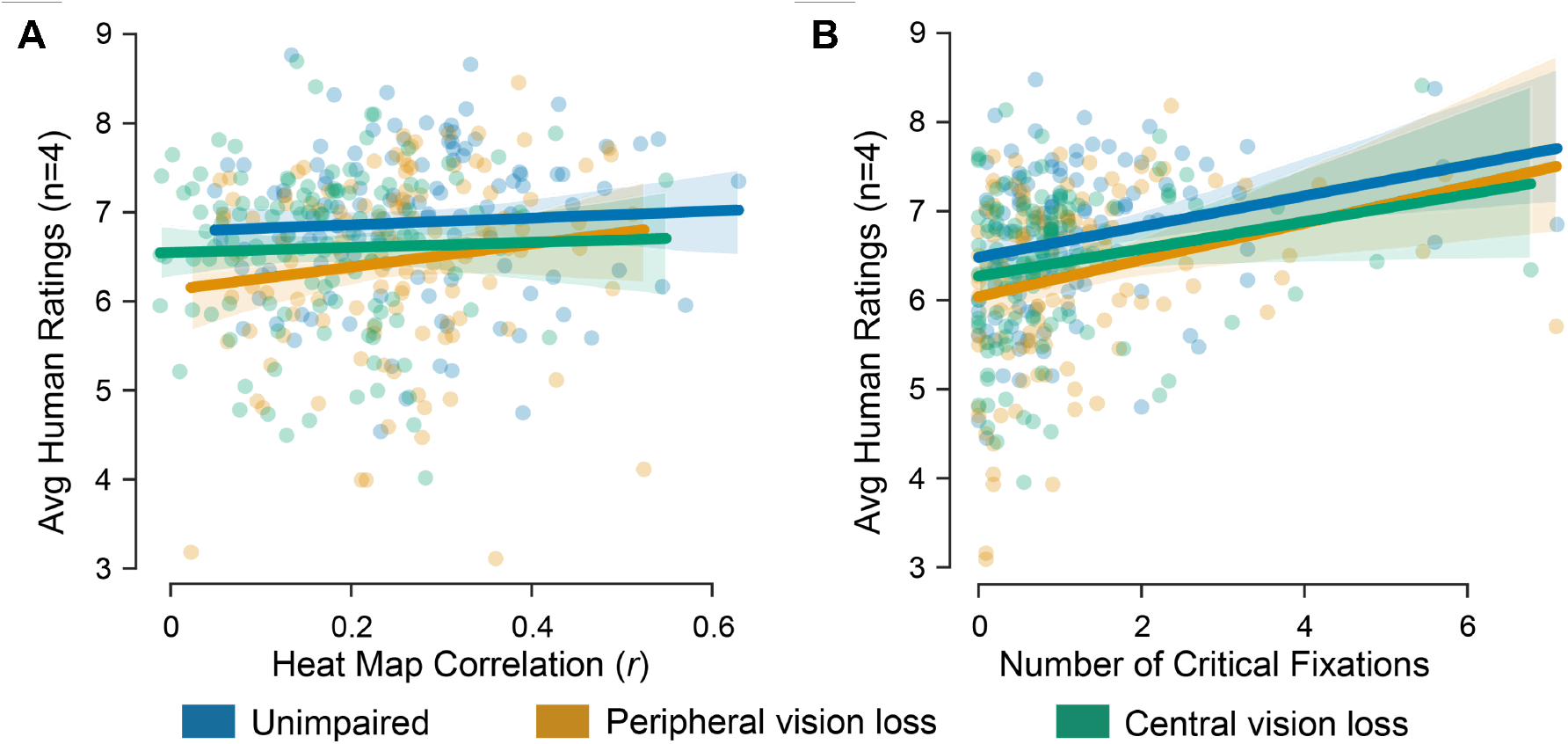
(A) Relationship between average human ratings of scene descriptions and fixation heat map correlations across all scenes. No significant association was observed, suggesting that global gaze distribution alone did not predict scene understanding. (B) Description ratings were significantly correlated with the number of critical fixations (i.e., fixations to annotated humans and critical objects), indicating that access to semantically meaningful regions of the scene contributed to higher-quality descriptions.

#### 3.3.1 Summary of Behavioral Measure Relationships

To explore how different behavioral and eye movement variables related to each other, we computed a correlation matrix using scene-level averages across all viewing conditions. The relationships between measures are shown in Figure 9. Description ratings were most strongly associated with the number of fixations to annotated humans and critical objects. In contrast, there was no reliable relationship between description ratings and global fixation similarity (i.e., heat map correlation). Interestingly, heat map correlation was negatively associated with first saccade amplitude, which itself was positively associated with first saccade latency. Both amplitude and latency were also positively correlated with overall scene viewing time.

**Figure 9.**
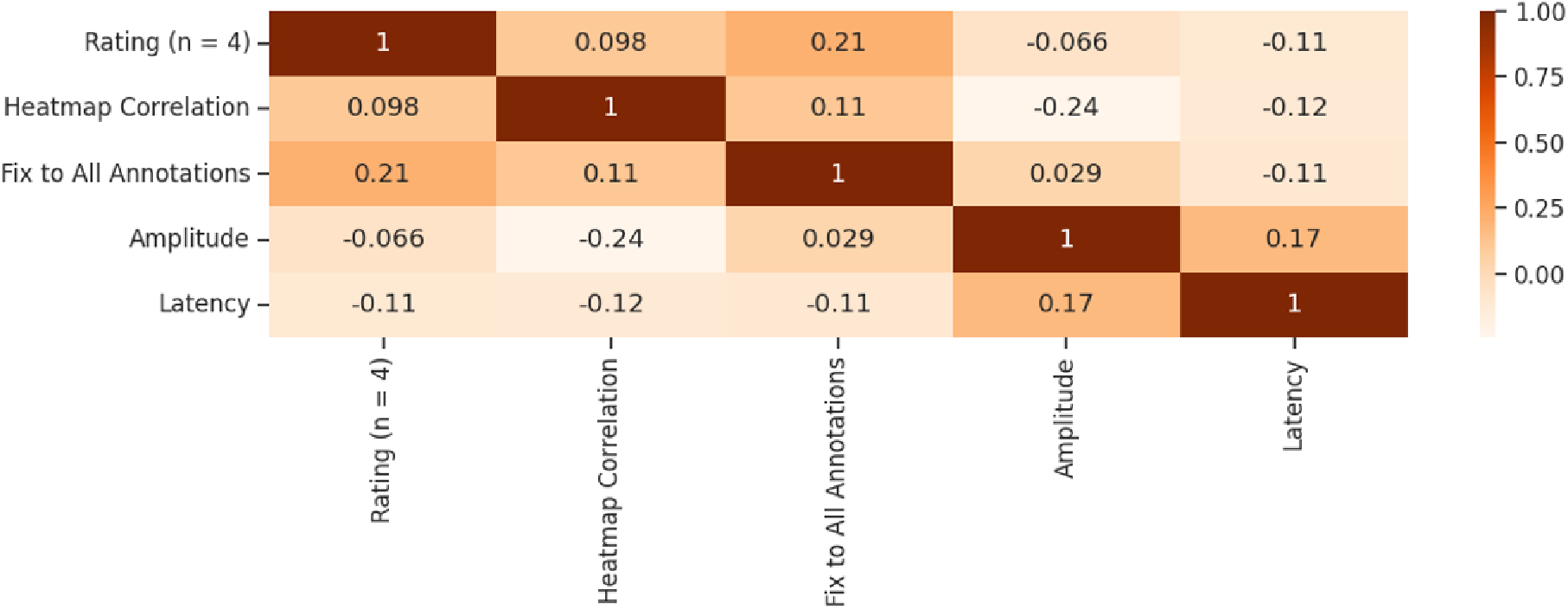
Correlation matrix of key behavioral and oculomotor measures.

## 4 Discussion

This study demonstrates that simulated vision loss (both central and peripheral) impairs rapid scene understanding and alters early eye movement behavior in complex, naturalistic environments. By combining gaze-contingent visual impairment with human-rated scene descriptions, we show that the type of vision loss produces distinct effects: CVL primarily disrupted where participants looked, while PVL strongly impacted their understanding of the scene. Importantly, comprehension suffered most in social scenes, where describing meaningful interactions requires integration of multiple scene elements. These findings suggest that rapid scene understanding relies on both foveal detail and peripheral context, and that different forms of visual impairment disrupt this process in complementary ways.

Our results extend prior work on scene categorization and simulated low vision (Loschky et al., 2019; Trouilloud et al., 2020) by using more ecologically valid stimuli and tasks. Descriptions from PVL and CVL participants were rated worse than controls, particularly for social interaction scenes (Fig. 4). This replicates and expands on prior evidence that peripheral vision supports rapid gist extraction (Thibaut et al., 2014; Larson and Loschky, 2009), and suggests that central vision is critical for resolving the social and semantic detail required to describe interactions. Notably, the neutral scenes (which often contained isolated objects or passive humans) produced smaller differences, likely because accurate description required only recognition rather than interpretation. Simulated vision loss also disrupted fixation behavior. CVL observers made larger first saccades (Fig. 5A) and had lower fixation heat map similarity to controls across all conditions (Fig. 6B– C). This contrasts with prior studies reporting smaller saccades in CVL patients with binocular scotoma during visual search (Vullings et al., 2022), or adaptive reductions in saccade range with training in simulations (Kwon et al., 2013; Vice et al., 2022). Our results suggest that under time pressure and without training, participants with central scotomas adopt a compensatory strategy of quickly fixating peripheral regions to avoid the blind spot. That PVL saccade amplitudes were indistinguishable from controls further supports the idea that CVL imposes stronger constraints on initial orienting.

Heat map similarity to controls did not predict description quality, even among control participants (Fig.8A), suggesting that meaningful scene understanding depends not on looking in the same place, but on extracting the right *information*. Supporting this, fixations to annotated humans and critical objects were consistently linked to higher-rated descriptions across all groups (Fig.8B), underscoring the importance of semantically informative fixations—particularly in social scenes.

Both PVL and CVL participants showed increased saccade latencies (Fig. 5B), consistent with prior studies of visual search and reading in patient populations and simulation paradigms (Ağaoğlu et al., 2022; Vice et al., 2022; Janssen and Verghese, 2016). Interestingly, CVL participants made faster and larger saccades than PVL participants, yet still produced higher-rated descriptions. One interpretation is that the central scotoma disrupted foveal detail but preserved access to scene context, whereas PVL observers struggled to localize targets and integrate spatial relationships due to a lack of peripheral preview. Supporting this view, saccade amplitude and latency were positively correlated in scenes with humans in the periphery (Fig. S8), particularly for CVL participants, suggesting that increased uncertainty delayed planning and triggered larger compensatory eye movements.

Taken together, our findings suggest that peripheral vision supports rapid orienting and context estimation, while central vision provides access to high-acuity details needed for interpretation. Both are necessary for functional scene understanding in the real world. Simulations provide an informative approximation of patient behavior, but future studies should replicate these findings in clinical populations. It remains an open question how much adaptation or training can mitigate the observed deficits with simulations (Vice et al., 2022; Nuthmann and Canas-Bajo, 2022; Ağaoğlu et al., 2022; McIlreavy et al., 2012), or how training gaze strategies can influence experience-dependent plasticity in patients with chronic vision loss (Janssen and Verghese, 2016). Longitudinal studies and eye movement–based interventions could reveal whether patients can learn to optimize gaze strategies under naturalistic conditions.

This work also highlights the value of naturalistic tasks, such as scene description, in assessing functional vision. Traditional clinical metrics like acuity and contrast sensitivity may not account for impairments in real-world understanding. Integrating eye tracking and semantically rich tasks could yield more sensitive and ecologically valid tools for evaluating patient outcomes and device efficacy. Furthermore, future work should explore how dynamic stimuli and active tasks, such as activities of daily living interact with vision loss. Our findings also emphasize the unique role of social information (i.e., faces, interactions, and people) in driving both visual behavior and semantic understanding. Functional assessments that ignore these dimensions may underestimate the real-world impact of vision loss.

Finally, while simulations are not a substitute for patient studies, they offer a safe and scalable way to prototype interventions and evaluate scene comprehension under controlled conditions. By anchoring behavioral performance in both gaze behavior and semantic interpretation, our findings help bridge the gap between sensory impairment and functional vision and point toward more ecologically valid tools for vision assessment and rehabilitation.

## AcknowledgmentS

The work was supported by a project from the Noyce Foundation. We thank Isabella Sarullo, Madeline Kaplan, Lucia Alem, Jin Hee Yang, Emma Bowen, and Jianna Wong for their assistance with data collection.

## Notes

Conflict of interest statement: The authors declare no competing financial interests.

### Competing Interest Statement

The authors have declared no competing interest.

## References

Ağaoğlu, M. N., Fung, W., and Chung, S. T. L. (2022). Oculomotor responses of the visual system to an artificial central scotoma may not represent genuine visuomotor adaptation. Journal of Vision, 22(10):17.

Bates, D., Mächler, M., Bolker, B., and Walker, S. (2015). Fitting Linear Mixed-Effects Models Using lme4. Journal of Statistical Software, 67(1).

Bradski, G. (2000). The OpenCV Library. Dr. Dobb’s Journal of Software Tools.

Costela, F. M., Saunders, D. R., Rose, D. J., Katjezovic, S., Reeves, S. M., and Woods, R. L. (2019). People With Central Vision Loss Have Difficulty Watching Videos. Investigative Opthalmology & Visual Science, 60(1):358.

Fletcher, D. C., Schuchard, R. A., and Renninger, L. W. (2012). Patient Awareness of Binocular Central Scotoma in Age-Related Macular Degeneration. Optometry and Vision Science, 89(9):1395–1398.

Geringswald, F. and Pollmann, S. (2015). Central and peripheral vision loss differentially affects contextual cueing in visual search. Journal of Experimental Psychology: Learning, Memory, and Cognition, 41(5):1485–1496.

Greene, M. R., Botros, A. P., Beck, D. M., and Fei-Fei, L. (2015). What you see is what you expect: rapid scene understanding benefits from prior experience. Attention, Perception, & Psychophysics, 77(4):1239–1251.

Guadron, L., Titchener, S. A., Abbott, C. J., Ayton, L. N., Van Opstal, J., Petoe, M. A., and Goossens, J. (2023). The Saccade Main Sequence in Patients With Retinitis Pigmentosa and Advanced Age-Related Macular Degeneration. Investigative Opthalmology & Visual Science, 64(3):1.

Hartong, D. T., Berson, E. L., and Dryja, T. P. (2006). Retinitis pigmentosa. The Lancet, 368(9549):1795–1809.

Janssen, C. P. and Verghese, P. (2016). Training eye movements for visual search in individuals with macular degeneration. Journal of Vision, 16(15):29.

Kasowski, J., Johnson, B. A., Neydavood, R., Akkaraju, A., and Beyeler, M. (2023). A systematic review of extended reality (XR) for understanding and augmenting vision loss. Journal of Vision, 23(5):5.

Kwon, M., Nandy, A., and Tjan, B. (2013). Rapid and Persistent Adaptability of Human Oculomotor Control in Response to Simulated Central Vision Loss. Current Biology, 23(17):1663–1669.

Larson, A. M. and Loschky, L. C. (2009). The contributions of central versus peripheral vision to scene gist recognition. Journal of Vision, 9(10):6–6.

Legge, G. E. and Chung, S. T. (2016). Low Vision and Plasticity: Implications for Rehabilitation. Annual Review of Vision Science, 2(1):321–343.

Lin, T.-Y., Maire, M., Belongie, S., Bourdev, L., Girshick, R., Hays, J., Perona, P., Ramanan, D., Zitnick, C. L., and Dollár, P. (2014). Microsoft COCO: Common Objects in Context. arXiv preprint.

Loschky, L. C., Szaffarczyk, S., Beugnet, C., Young, M. E., and Boucart, M. (2019). The contributions of central and peripheral vision to scene-gist recognition with a 180° visual field. Journal of Vision, 19(5):15.

McIlreavy, L., Fiser, J., and Bex, P. J. (2012). Impact of Simulated Central Scotomas on Visual Search in Natural Scenes. Optometry and Vision Science, 89(9):1385–1394.

Nguyen, D., Trieschnigg, D., and Theune, M. (2014). Using Crowdsourcing to Investigate Perception of Narrative Similarity. In Proceedings of the 23rd ACM International Conference on Conference on Information and Knowledge Management, pages 321–330, Shanghai China. ACM.

Nuthmann, A. (2014). How do the regions of the visual field contribute to object search in real-world scenes? Evidence from eye movements. Journal of Experimental Psychology: Human Perception and Performance, 40(1):342–360.

Nuthmann, A. and Canas-Bajo, T. (2022). Visual search in naturalistic scenes from foveal to peripheral vision: A comparison between dynamic and static displays. Journal of Vision, 22(1):10.

Peirce, J., Gray, J. R., Simpson, S., MacAskill, M., Höchenberger, R., Sogo, H., Kastman, E., and Lindeløv, J. K. (2019). PsychoPy2: Experiments in behavior made easy. Behavior Research Methods, 51(1):195–203.

Peli, E., Goldstein, R., and Jung, J.-H. (2023). The Invisibility of Scotomas I: The Carving Hypothesis. Optometry and Vision Science, 100(8):515–529.

Peyrin, C., Ramanöel, S., Roux-Sibilon, A., Chokron, S., and Hera, R. (2017). Scene perception in age-related macular degeneration: Effect of spatial frequencies and contrast in residual vision. Vision Research, 130:36–47.

Seiple, W., Rosen, R. B., and Garcia, P. M. (2013). Abnormal Fixation in Individuals With Age-Related Macular Degeneration When Viewing an Image of a Face. Optometry and Vision Science, 90(1):45–56.

Shintani, K., Shechtman, D. L., and Gurwood, A. S. (2009). Review and update: Current treatment trends for patients with retinitis pigmentosa. Optometry - Journal of the American Optometric Association, 80(7):384–401.

Skalski, P. (2019). Make Sense.

Thibaut, M., Tran, T. H. C., Szaffarczyk, S., and Boucart, M. (2014). The contribution of central and peripheral vision in scene categorization: A study on people with central vision loss. Vision Research, 98:46–53.

Tran, T. H. C., Rambaud, C., Despretz, P., and Boucart, M. (2010). Scene Perception in Age-Related Macular Degeneration. Investigative Opthalmology & Visual Science, 51(12):6868.

Trouilloud, A., Kauffmann, L., Roux-Sibilon, A., Rossel, P., Boucart, M., Mermillod, M., and Peyrin, C. (2020). Rapid scene categorization: From coarse peripheral vision to fine central vision. Vision Research, 170:60–72.

Tsank, Y. and Eckstein, M. P. (2017). Domain Specificity of Oculomotor Learning after Changes in Sensory Processing. The Journal of Neuroscience, 37(47):11469–11484.

Van Der Stigchel, S., Bethlehem, R. A. I., Klein, B. P., Berendschot, T. T. J. M., Nijboer, T. C. W., and Dumoulin, S. O. (2013). Macular degeneration affects eye movement behavior during visual search. Frontiers in Psychology, 4.

Verghese, P., Vullings, C., and Shanidze, N. (2021). Eye Movements in Macular Degeneration. Annual Review of Vision Science, 7(1):773–791.

Vice, J. E., Biles, M. K., Maniglia, M., and Visscher, K. M. (2022). Oculomotor changes following learned use of an eccentric retinal locus. Vision Research, 201:108126.

Vullings, C., Lively, Z., and Verghese, P. (2022). Saccades during visual search in macular degeneration. Vision Research, 201:108113.

Wang, Y., Tao, S., Xie, N., Yang, H., Baldwin, T., and Verspoor, K. (2023). Collective Human Opinions in Semantic Textual Similarity. Transactions of the Association for Computational Linguistics, 11:997–1013.

Yu, H. and Kwon, M. (2023). Altered Eye Movements During Reading With Simulated Central and Peripheral Visual Field Defects. Investigative Opthalmology & Visual Science, 64(13):21.

