## Supplemental Figures for "Distinct Roles of Central and Peripheral Vision in Rapid Scene Understanding"

### Supplemental Material

#### Timing Validation for Gaze-Contingent Display

To estimate the delay between eye position updates and stimulus rendering, we inserted timestamped logging commands into the experiment's Python code. For 10 randomly selected scenes, we recorded the time elapsed between receiving gaze coordinates from the eye tracker and rendering the updated frame with the appropriate scotoma filter. The average delay was $6.89 \pm 1.21\,\mathrm{ms}$ across all tested trials (Figure S1).

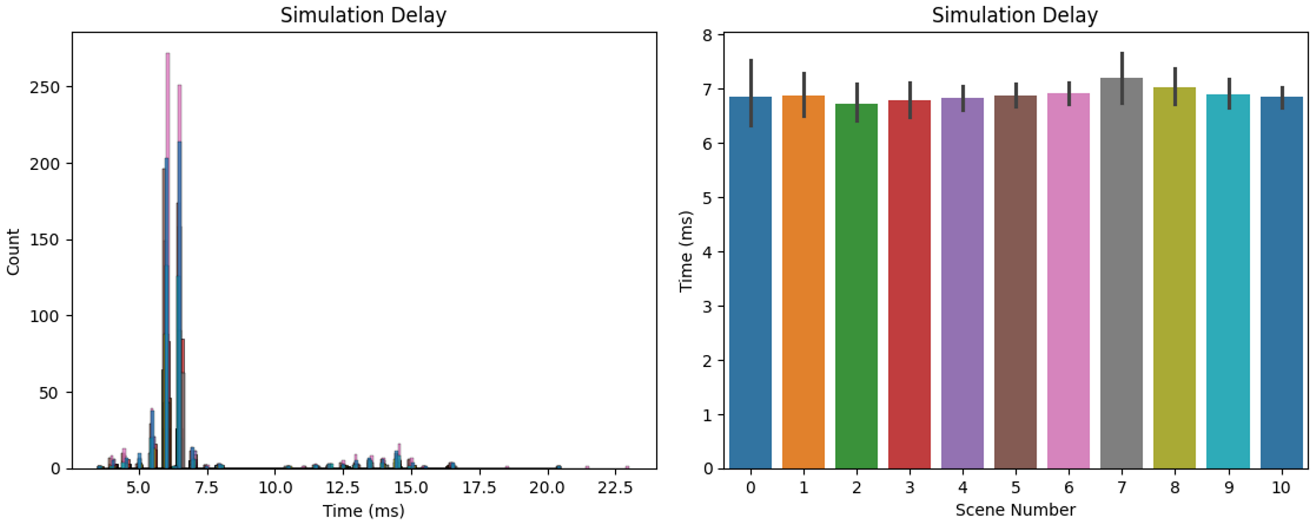


Figure S1: Timing validation of gaze-contingent display updates. The histogram (left) shows the distribution of delays across trials and scenes. The right panel shows the mean and standard deviation per trial for each scene.

#### Human Rater Reliability and Rating Distributions

To assess variability and agreement in human semantic similarity judgments, we analyzed the distribution of ratings, pairwise inter-rater reliability, and rating correlations between all four raters.

Figure S2A shows the distribution of ratings assigned by each rater across all scene descriptions. A Shapiro–Wilk test indicated that the average ratings were not normally distributed ($W = 0.98$, $p < .001$). The mean rating across all descriptions was 6.44 (SEM = 0.02), and both the mode and median were 6.5.

Figure S2B shows pairwise inter-rater reliability (IRR) scores between raters, calculated as the proportion of agreement in ordinal ranking across all rated items. IRR values ranged from 0.042 (Rater 1 and Rater 4) to 0.23 (Rater 2 and Rater 4), indicating modest agreement in relative similarity judgments.

Figure S2C presents a Pearson correlation matrix across all rater pairs. All correlations were statistically significant ($p < .001$), with coefficients ranging from $r = 0.32$ (Rater 1 and Rater 4) to $r = 0.74$ (Rater 2 and Rater 3). These analyses demonstrate moderate consistency across raters and support the use of average ratings in subsequent analyses.

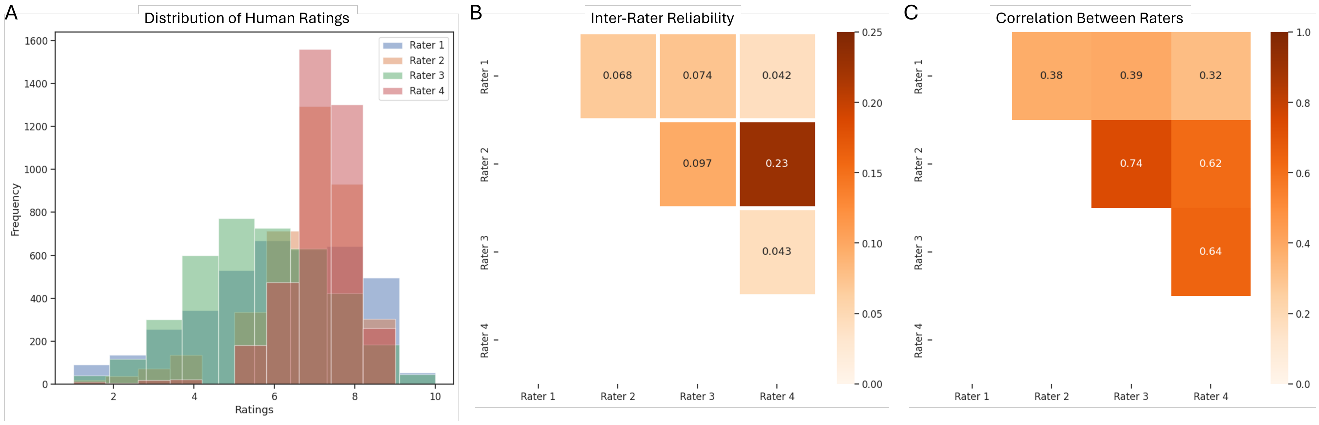


Figure S2: **A)** Distribution of semantic similarity ratings by each of the four human raters across all 3,840 scene descriptions. **B)** Pairwise inter-rater reliability (IRR) values based on ordinal agreement across all descriptions. Values indicate modest agreement, especially between Rater 2 and the other raters. **C)** Pearson correlation matrix between all human raters. All pairwise correlations were significant at $p < .001$, with strongest agreement between Rater 2 and Rater 3.

#### Stability of Scene Description Performance Over Time

To assess whether participants improved their descriptions over time, we compared semantic similarity scores from the first and last experimental blocks, separately for each viewing condition. Figure S3 shows the average human ratings (top) and GPT-4 cosine similarity scores (bottom) across blocks. A two-way ANOVA with viewing condition and block number as factors found no significant effect of block number on similarity scores ($F(1, 6) = 1.27$, $p = .303$), indicating no evidence of learning across the experiment.

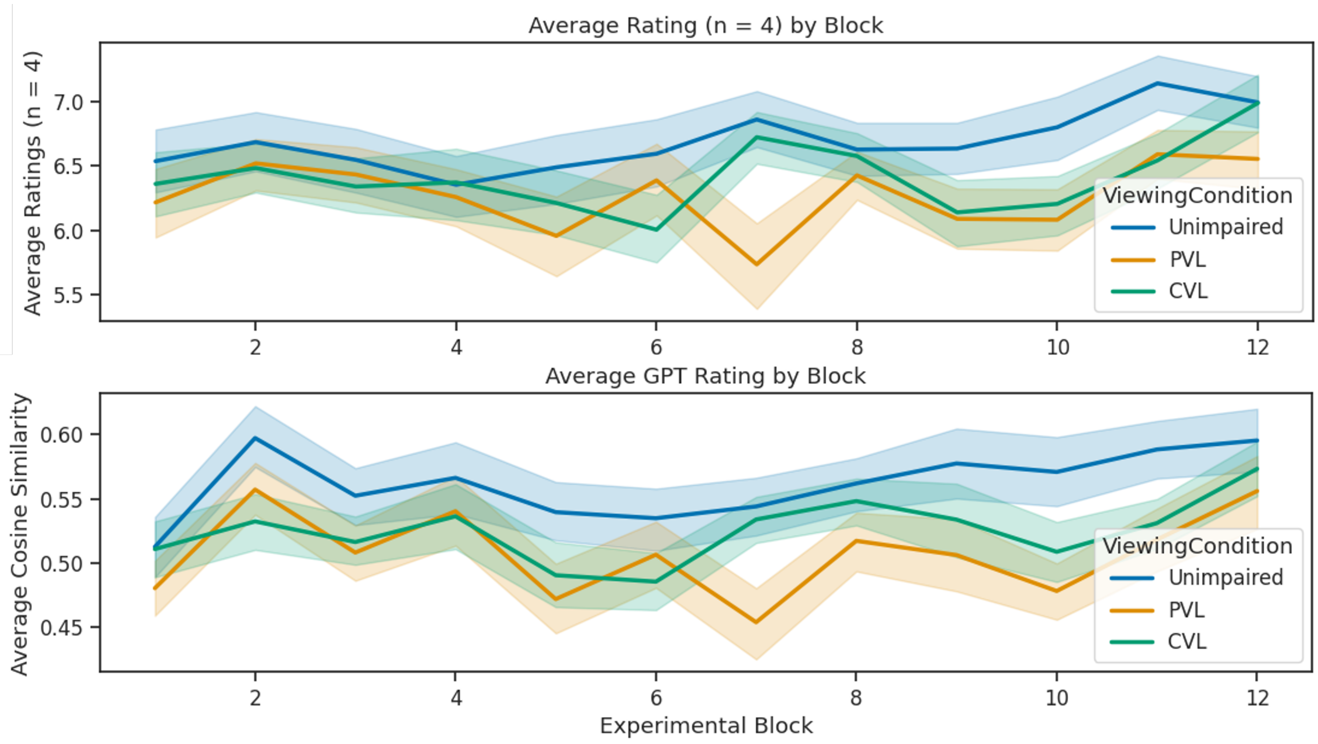


Figure S3: Average semantic similarity ratings by block number and viewing condition. Top: human ratings; bottom: GPT-4 cosine similarity. No significant learning effects were observed across blocks.

#### Example Heat Maps and Stimulus Annotations

To illustrate the structure and variability of our stimulus set, this section provides representative examples of group-level fixation heat maps and annotated scenes.

To visualize group-level gaze behavior, fixation locations were transformed into fixation heat maps for each scene. Heat maps were generated by randomly sampling groups of 4 participants from each viewing condition (control, PVL, CVL) as well as from a separate "control comparability group." Fixation locations were convolved with a Gaussian kernel ($39 \times 39$ pixels, approximately $1 \times 1°$) to estimate spatial fixation density. Red areas indicate regions of high fixation density. Figures S4, S5, and S6 shows example heat maps for one scene across the four groups.

Each of the 120 video scenes was manually annotated to identify the faces and bodies of humans or animals, as well as critical task-relevant objects. Annotations were created using Make Sense (Skalski, 2019) and saved in JSON format. A critical object was defined as the object (or group of objects) most relevant for accurately describing the scene. Annotations were also categorized as either central or peripheral, based on whether they appeared beyond 5 degrees of visual angle from the initial fixation point. Of the 120 scenes, 61 included people in the periphery (45 social, 16 neutral), and 33 were labeled as containing peripheral critical objects (21 social, 12 neutral). Examples of annotated frames from both social interaction and neutral scenes are shown in Figures S7.

These examples illustrate how viewing condition influenced fixation behavior relative to the spatial distribution of social and task-relevant content.

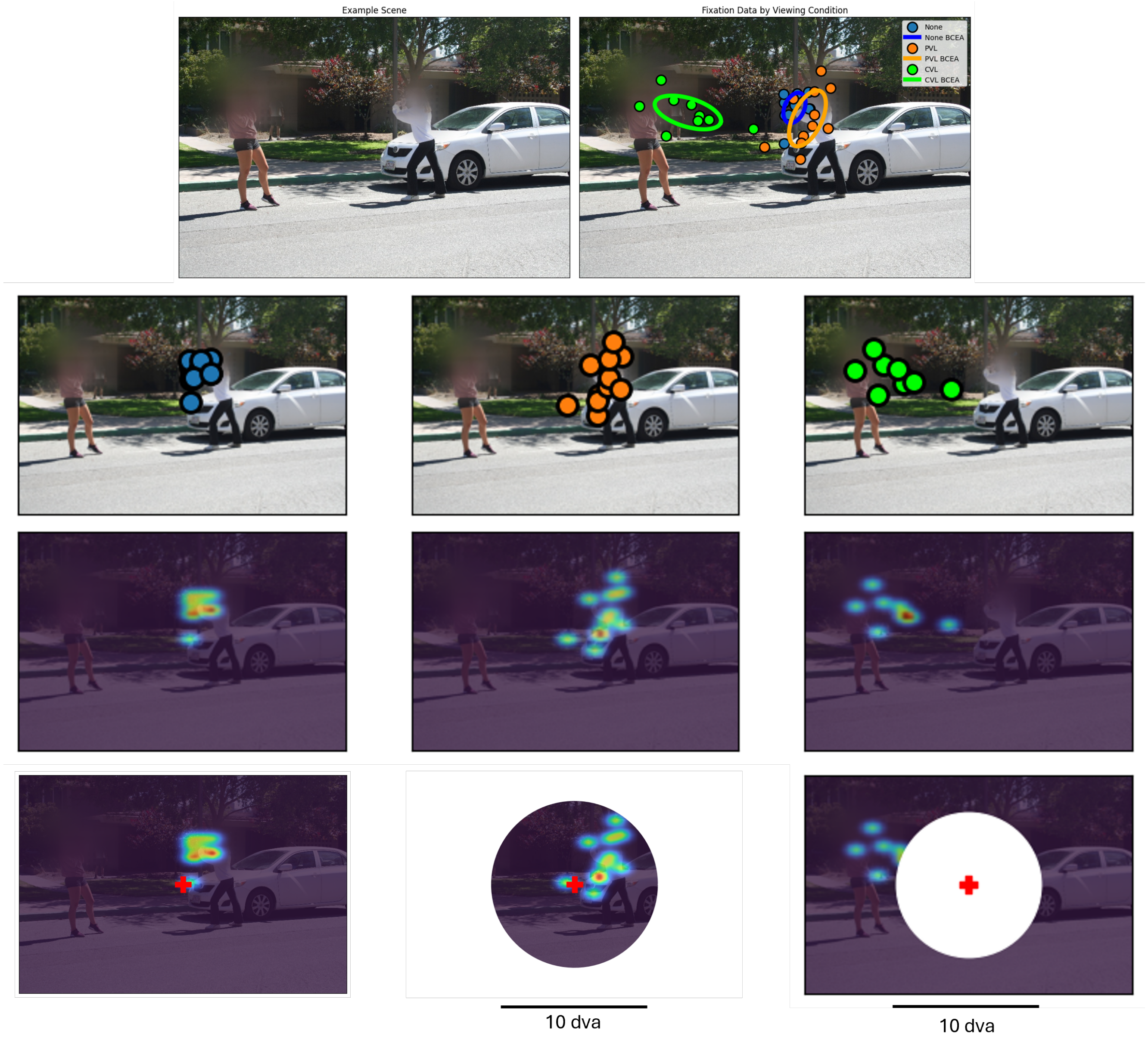


Figure S4: Example scene with raw fixation data and heat maps by viewing condition: (top left) original scene, (top right) scene with fixation data based on viewing condition, (second row) fixation data for controls, PVL, and CVL, (third row) convolving the fixation data with a Gaussian filter, (fourth row) heat map data with representation of experimental viewing conditions at stimulus onset. For example trial of the actual experiment, refer to Figure 2. Faces in example social interaction scenes are blurred for sharing of this manuscript.

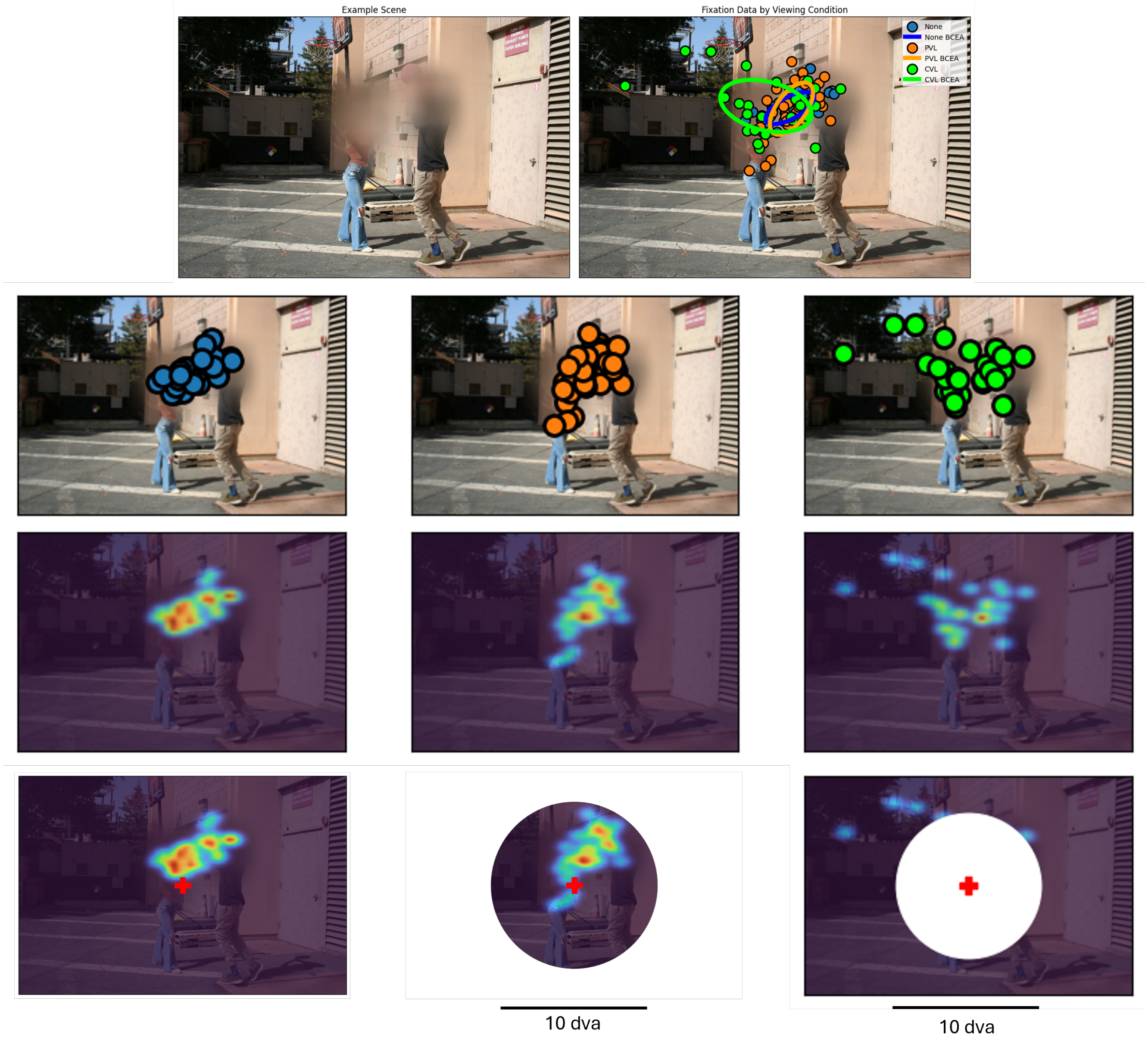


Figure S5: Example social interaction scene of people arranged centrally with raw fixation data and heat maps by viewing condition: (top left) original scene, (top right) scene with fixation data based on viewing condition, (second row) fixation data for controls, PVL, and CVL, (third row) convolving the fixation data with a Gaussian filter, (fourth row) heat map data with representation of experimental viewing conditions at stimulus onset. For example trial of the actual experiment, refer to Figure 2. Faces in example social interaction scenes are blurred for sharing of this manuscript.

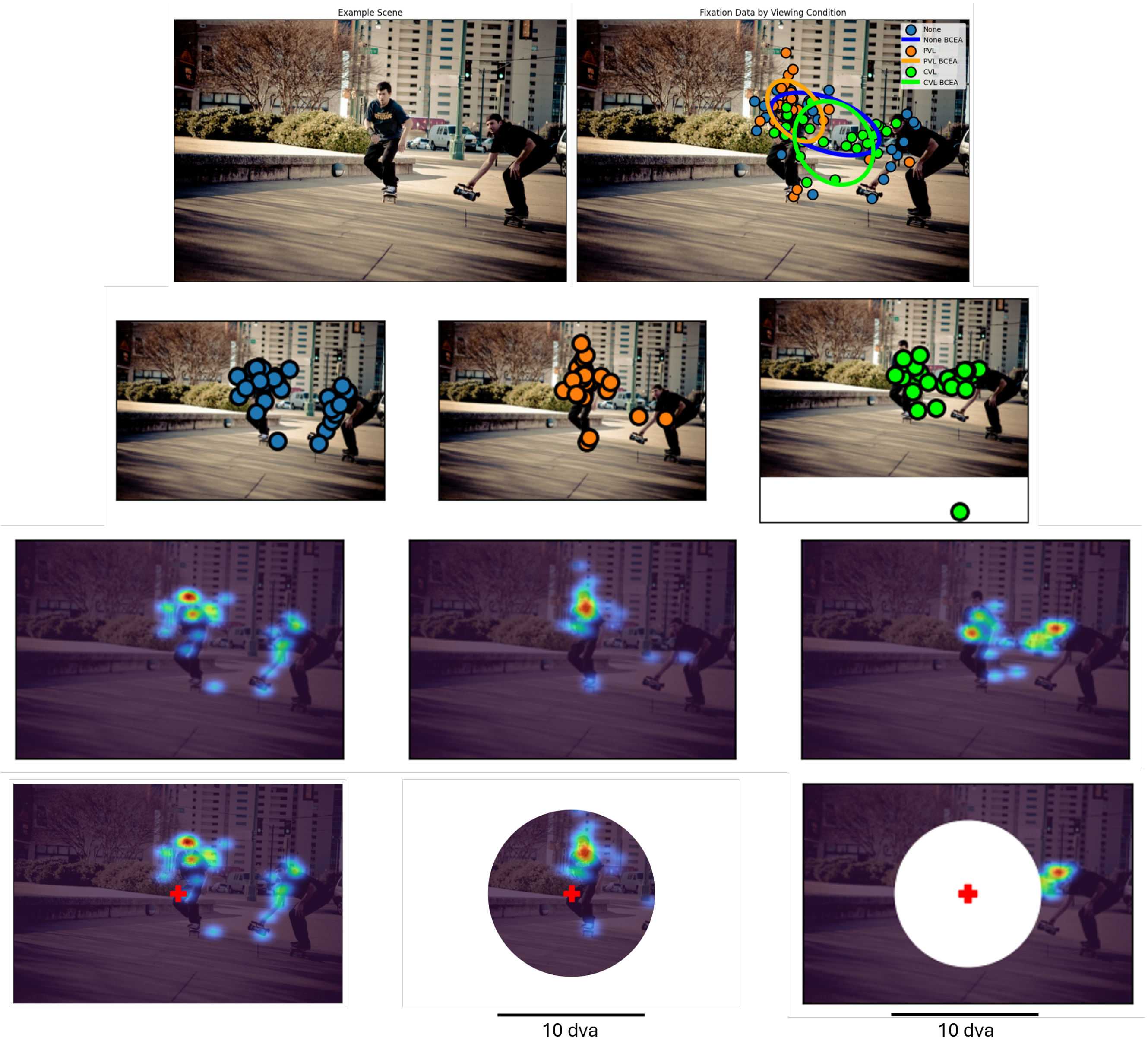


Figure S6: Example neutral scene with a person arranged peripherally with raw fixation data and heat maps by viewing condition: (top left) original scene, (top right) scene with fixation data based on viewing condition, (second row) fixation data for controls, PVL, and CVL, (third row) convolving the fixation data with a Gaussian filter, (fourth row) heat map data with representation of experimental viewing conditions at stimulus onset. For example trial of the actual experiment, refer to Figure 2.

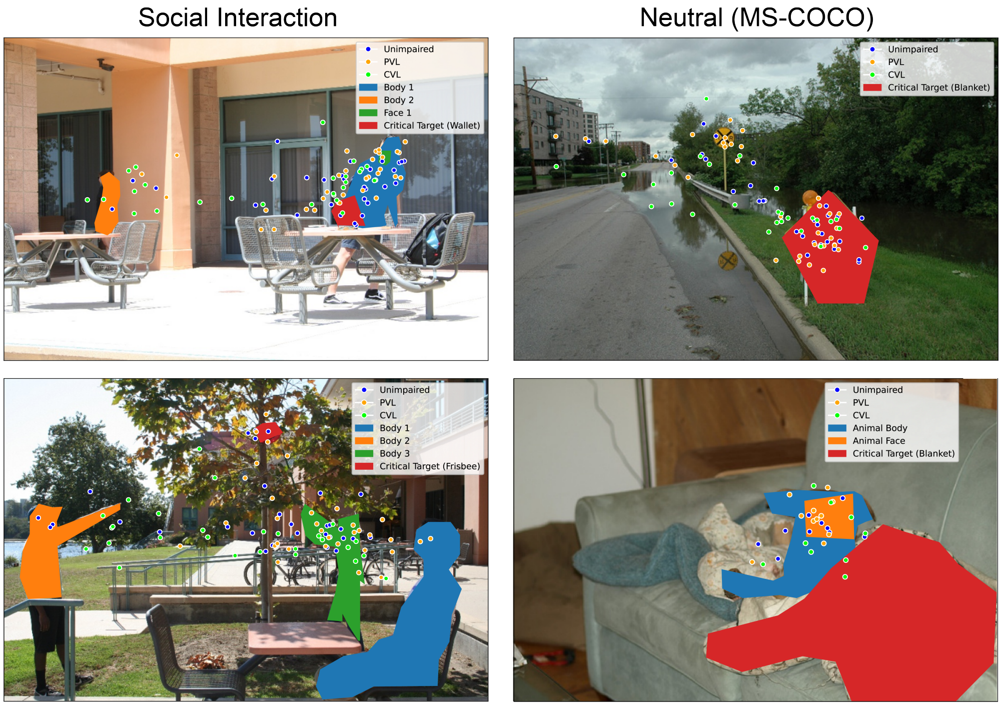


Figure S7: Example scenes with fixation locations and labels for humans and critical objects, for social scenes (left) and neutral scenes (right).

#### Saccade Metrics for Scenes With and Without Peripheral Humans

To further examine the role of peripheral social cues, we analyzed the relationship between first saccade amplitude and latency, stratified by scene type (social interactions vs. neutral), presence of humans in the periphery, and viewing condition (Fig. S8). This analysis supports the idea that peripheral social content increases oculomotor uncertainty and planning time, particularly under central vision loss.

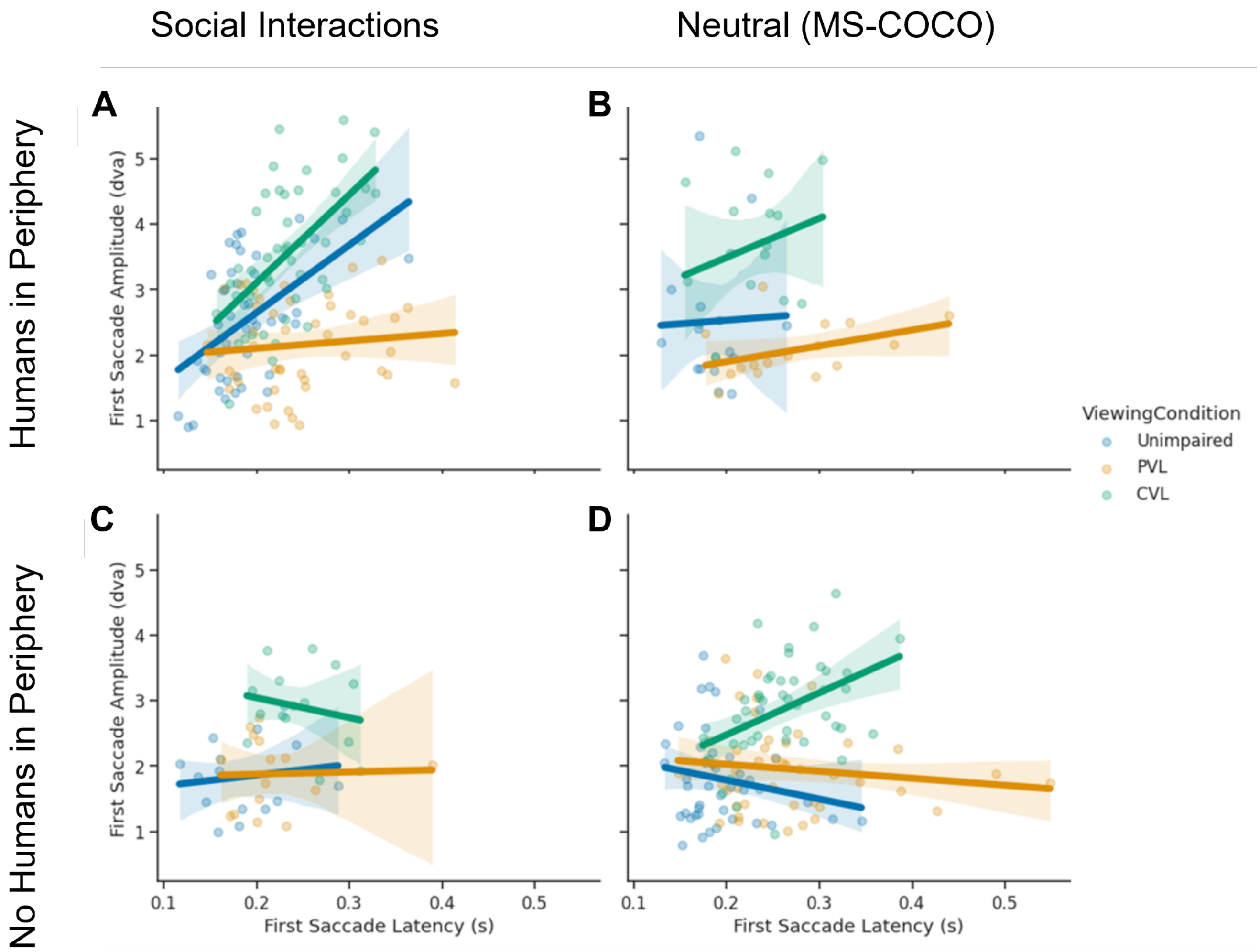


Figure S8: Relationship between first saccade amplitude and latency as a function of scene type, peripheral human presence, and viewing condition. In scenes with humans in the periphery (Panels A and B), saccade amplitude and latency were positively correlated, especially for CVL participants, suggesting delayed saccade planning and compensatory eye movements. This relationship was weaker or absent in scenes without peripheral humans (Panels C and D).
